# A chromatin phase transition protects mitotic chromosomes against microtubule perforation

**DOI:** 10.1101/2021.07.05.450834

**Authors:** Maximilian W. G. Schneider, Bryan A. Gibson, Shotaro Otsuka, Maximilian F.D. Spicer, Mina Petrovic, Claudia Blaukopf, Christoph C. H. Langer, Lynda K. Doolittle, Michael K. Rosen, Daniel W. Gerlich

## Abstract

Dividing eukaryotic cells package extremely long chromosomal DNA molecules into discrete bodies to enable microtubule-mediated transport of one genome copy to each of the newly forming daughter cells^1–3^. Assembly of mitotic chromosomes involves DNA looping by condensin^4–8^ and chromatin compaction by global histone deacetylation^9–13^. While condensin confers mechanical resistance towards spindle pulling forces^14–16^, it is not known how histone deacetylation affects material properties and segregation mechanics of mitotic chromosomes. Here, we show how global histone deacetylation at the onset of mitosis induces a chromatin-intrinsic phase transition that endows chromosomes with specific characteristics necessary for their precise movement during cellular division. Deacetylation-mediated compaction of chromatin forms a structure dense in negative charge and allows mitotic chromosomes to resist perforation by microtubules as they are pushed to the metaphase plate. Hyperacetylated mitotic chromosomes lack a defined surface boundary, are frequently perforated by microtubules, and are prone to missegregation. Our study highlights the different contributions of DNA loop formation and chromatin-intrinsic phase separation to genome segregation in dividing cells.

## Main text

The material properties of individual cell components play a key role in the dynamic organization and reorganization of cellular structure. In mitotic vertebrate cells, chromosomes must acquire material properties that enable microtubules to move them, first to the spindle center during prometaphase and then to the spindle poles during anaphase^1,17^. Pulling forces are generated by microtubules that attach to small chromosomal regions termed kinetochores^18^, whereas pushing forces, known as polar ejection forces, are produced by microtubules contacting chromosome arms^19–23^. Condensin confers mitotic chromosomes with the mechanical stability required to withstand pulling forces^14–16^, but it remains unclear how chromosomes resist pushing forces. These material properties must also provide chromosome arms with resistance to penetration by polymerizing microtubule tips, since entanglements occurring due to growth of microtubules into the mass of chromatin would lead to impaired segregation.

### Acetylation-regulated chromatin compaction prevents microtubule perforation

To investigate the relationship between the material properties of chromosomes and the forces exerted upon them by the mitotic spindle, we first sought to uncouple the mechanisms responsible for providing resistance to pushing and pulling forces. Given its well described role in resisting pulling forces^14–16^, we first investigated the effect of condensin depletion on the morphology and movement of mitotic chromosomes in live HeLa cells. To deplete condensin, we modified all endogenous alleles of its essential SMC4 subunit with a C-terminal auxin-inducible degron (mAID) and HaloTag for visualization and added the auxin analog 5-PhIAA^24^ 2.5 h prior to mitotic entry to induce efficient degradation (Extended Data Fig. 1a-c). We visualized the spindle with silicon-rhodamine (SiR)-tubulin^25^ and stained DNA with Hoechst to determine the position of chromosomes. In condensin-depleted cells, we observed an unstructured mass of compact chromatin forming a plate between the spindle poles (Fig. 1a, b), indicating that polar ejection forces efficiently push bulk chromatin towards the spindle center. Immunofluorescence staining of the kinetochore marker CENP-A and the spindle pole component pericentrin further showed that many kinetochores were detached from the bulk mass of chromatin and displaced towards the spindle poles, whereas in control cells all kinetochores were closely linked with chromosomes at the metaphase plate (Extended Data Fig. 1d). The movement of loosely connected kinetochores towards the spindle poles confirms that condensin is required for chromosomes to resist pulling forces. Thus, condensin depletion allows the mechanisms responsible for the resistance to pushing and pulling forces to be uncoupled.

**Figure 1.**
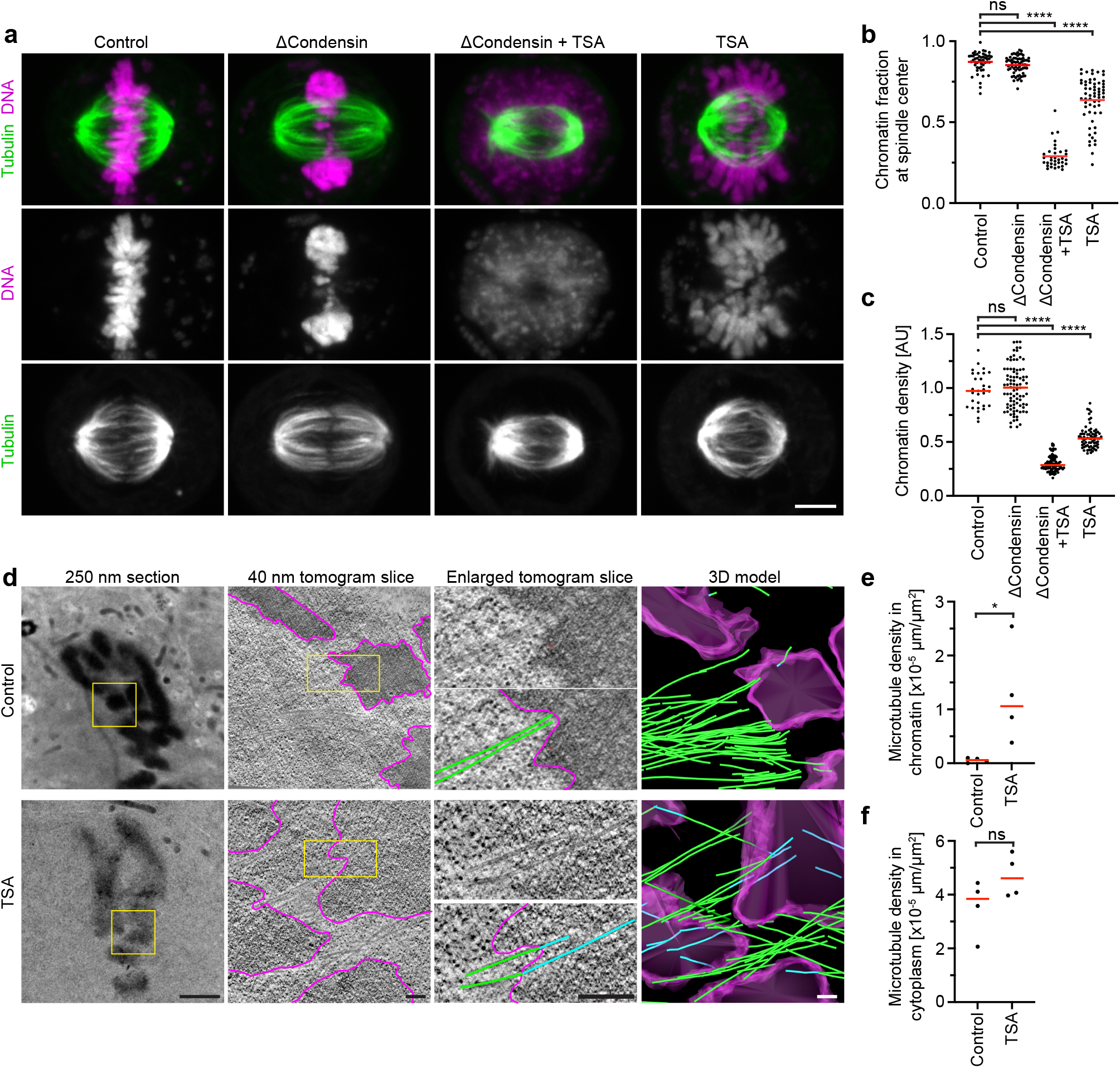
Acetylation-regulated chromatin compaction prevents microtubule perforation in mitosis. **a**, Contribution of condensin and histone deacetylases to mitotic chromosome compaction and congression to the spindle center. HeLa cells with homozygously mAID-tagged SMC4 were treated with 5-PhIAA to deplete condensin (ΔCondensin) or with TSA to suppress mitotic histone deacetylation as indicated. Live cell images with microtubules stained by SiR-tubulin; DNA stained by Hoechst. Projection of 5 Z-sections. **b**, Quantification of chromosome congression by the fraction of chromatin localizing to the central spindle region. *n*=51 cells for control, *n*=65 for ΔCondensin, *n*=34 for ΔCondensin+TSA, *n*=61 for TSA. Bars indicate mean; significance was tested by an unpaired, two-tailed *t*-test (ΔCondensin, *P*=0.06) or a two-tailed Mann-Whitney test (ΔCondensin+TSA, *P*<10^−15^); TSA, *P*<10^−15^). **c**, Quantification of chromatin density in cells treated as in **a**. *n*=31 cells for control, *n*=89 for ΔCondensin, *n*=99 for ΔCondensin+TSA, *n*=74 for TSA. Bars indicate mean; significance was tested by an unpaired, two-tailed *t*-test (ΔCondensin, *P*=0.816) or a Kolmogorov-Smirnov test (ΔCondensin+TSA, *P*<10^−15^; TSA, *P*<10^−15^). **d**, Electron tomography of wildtype prometaphase HeLa cells in the absence or presence of TSA. **e, f**, Quantification of microtubule density in chromatin and cytoplasmic regions as shown in **d**. *n*=4 tomograms from 3 cells for each condition. Bars indicate mean; significances were tested by a two-tailed Mann-Whitney test (**e,** *P*=0.029; *P*=0.343). Scale bars, **a**, 5 μm, **d**, 250 nm section, 2 μm, tomogram slices, 200 nm.

The observation that bulk chromatin remains responsive to polar ejection forces following depletion of condensin suggests that under these conditions resistance to pushing forces might arise from nucleosome-mediated interactions in the chromatin fiber. Nucleosomal interactions are thought to increase when histones are deacetylated during mitotic entry (Extended Data Fig. 2a, b), contributing to global chromatin compaction^9–11,26^. To investigate how acetylation affects the movement of bulk chromatin, we treated condensin-depleted cells with the pan histone deacetylase inhibitor trichostatin A (TSA)^27^ to induce broad hyperacetylation of histone lysine residues (Extended Data Fig. 2c, d). The hyperacetylated chromatin of condensin-depleted mitotic cells did not compact or concentrate at the spindle center but instead dispersed throughout the cytoplasm, such that spindle pole regions were no longer clear of chromatin (Fig. 1a, b; Extended Data Fig. 1d; 2e, f). A complementary approach to induce histone hyperacetylation by overexpressing the histone acetyltransferase p300^28^ resulted in consistent chromatin decompaction and congression phenotypes (Extended Data Fig. 3). Thus, histone deacetylation appears to be important for spindle-mediated pushing of bulk chromatin towards the spindle center.

The chromatin decompaction induced by TSA in condensin-depleted cells appeared much more pronounced than previously observed in wildtype cells^9,10^. We hypothesized that this might be due to condensin-mediated linkages that counteract the dispersion of chromatin fibers. To investigate this, we analyzed how TSA affects chromatin density in the presence or absence of condensin. In condensin-depleted mitotic cells, TSA reduced chromatin density to 29% compared to cells that were not treated with TSA, whereas in the presence of condensin it reduced chromatin density only to 53% (Fig. 1a, c). Condensin depletion alone did not reduce mitotic chromatin density (Fig. 1a, c), consistent with previous observations^14,29,30^. Thus, histone deacetylation is necessary and sufficient for complete compaction of mitotic chromatin even in the absence of condensin. In contrast, condensin is neither necessary nor sufficient for complete chromatin compaction during mitosis, yet it can concentrate chromatin to some extent even when histones are hyperacetylated. These observations support a model describing mitotic chromosomes as a hydrogel^31–34^, in which a flexible chromatin fiber is linked by condensin to form a network, wherein the degree of compaction is controlled by acetylation-regulated interactions between nucleosomes, resulting in changes in fiber solubility.

The hydrogel model explains how condensin linkages confer resistance to pulling forces^14–16^. Histone deacetylation appears to be dispensable for resistance against pulling forces^35^, yet a potential role in facilitating microtubule-based pushing has not been investigated. We hypothesized that resistance to pushing forces depends first on the ability to prevent microtubules penetrating chromosomes, which might require compaction of a swollen interphase chromatin hydrogel via histone deacetylation during mitotic entry. To test how acetylation affects access of microtubules to chromosomes, we performed electron tomography of mitotic HeLa cells. Chromosomes of unperturbed cells appeared as homogeneously compacted bodies with a sharp surface boundary, and 3D segmentation showed that they were almost never penetrated by microtubules (Fig. 1d, e). In contrast, chromosomes of TSA-treated cells appeared less compact, particularly towards the periphery, and microtubules grew extensively through the chromatin (Fig. 1d-f). Thus, active histone deacetylases are required to keep microtubules out of chromosome bodies, providing a basis for resistance towards pushing forces.

Microtubule perforation into mitotic chromosomes is expected to cause entanglements between chromatin fiber loops and spindle microtubules that impair chromosome segregation. To investigate how TSA-induced hyperacetylation affects chromosome movements in mitosis, we recorded time-lapse movies of HeLa cells expressing H2B-mCherry. TSA severely delayed chromosome congression and initiation of anaphase and caused a high incidence of lagging chromosomes (Extended Data Fig. 1e-g). Active histone deacetylases are thus essential for faithful chromosome segregation.

### Acetylation controls chromatin solubility in cytoplasm

To elucidate the mechanism underlying microtubule exclusion from mitotic chromosomes, we investigated how histone acetylation affects the material properties of chromatin. Both intact mitotic chromosomes^9,11^ and intrinsic chromatin condensates formed from small fragments of chromatin^13^ exhibit structural sensitivity to histone hyperacetylation. We reasoned, in accordance with principles from polymer chemistry^36^, that the hydrogel material of mitotic chromosomes that resists microtubule perforation might arise as a consequence of intrinsic chromatin phase separation at the very long length scale of human chromosomes.

To investigate the phase separation behavior of mitotic chromatin at smaller length scales, we fragmented chromosomes in live mitotic HeLa cells by microinjecting the restriction enzyme AluI. Shortly after microinjection of AluI, chromosomes lost their elongated shape, forming round condensates that fused to one another, indicating a liquid state (Fig. 2a). Remarkably, the digestion of chromatin did not decrease chromatin density (Fig. 2b), demonstrating that integrity of the chromatin fiber is not required for full compaction of mitotic chromatin. At macroscopic scales, phase-separating systems become less dynamic when composed of long molecules due to steric effects and length-dependent changes in valency. Consistent with this intrinsic behavior, without AluI digestion H2B-mCherry fluorescence recovers very little after photobleaching, whereas mitotic chromatin condensates that form following AluI digestion recover rapidly from photobleaching (Fig. 2c, d). Thus, mitotic chromatin is insoluble in cytoplasm, and when the long-range constraints of the fiber are eliminated, the short-range dynamics manifest in liquid-like behavior.

**Figure 2.**
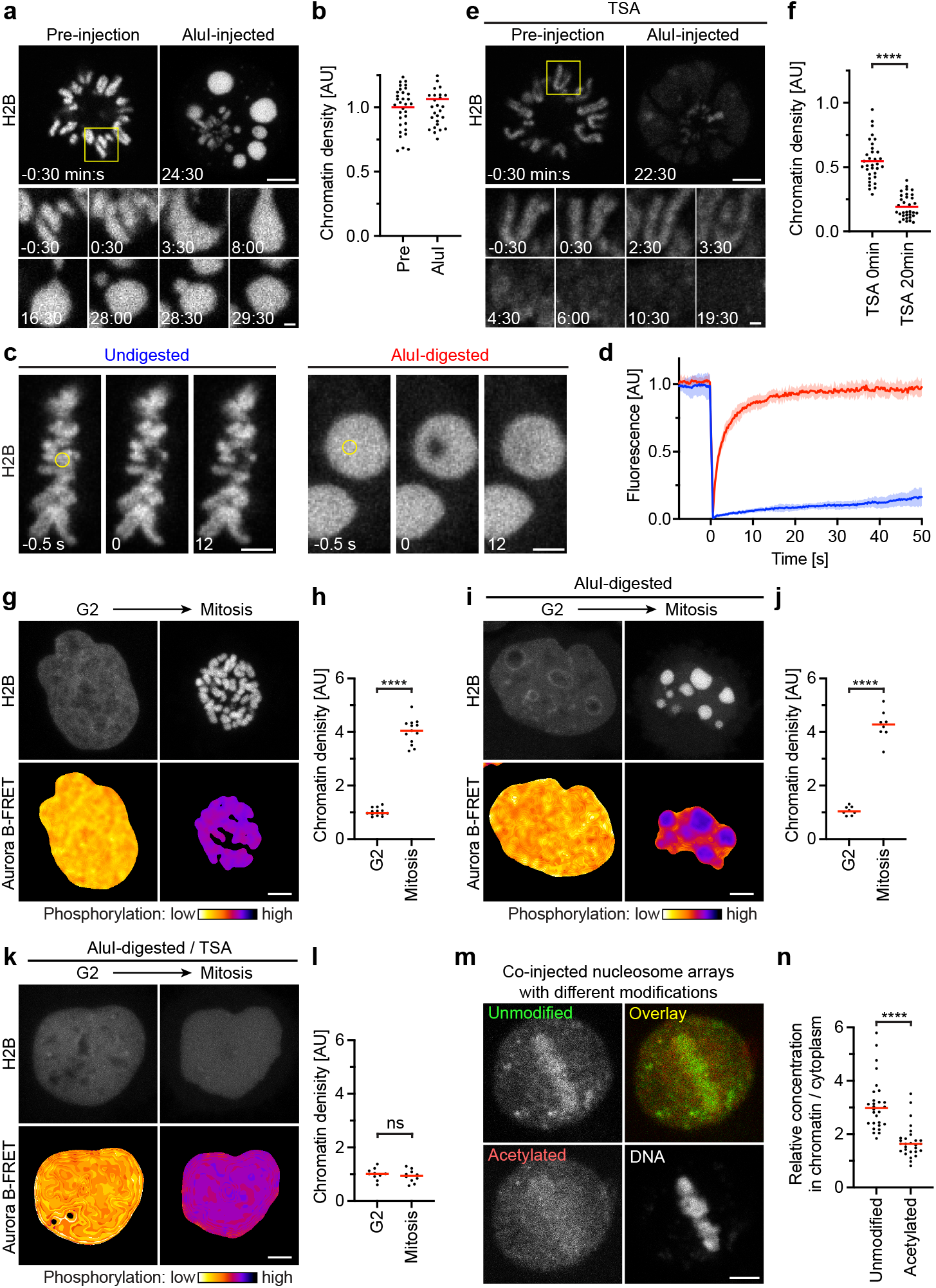
Acetylation regulates chromatin solubility in mitotic cytoplasm. **a**, Chromosome fragmentation in live mitotic HeLa cell by AluI injection during time-lapse microscopy. Chromatin was visualized with H2B-mCherry. AluI was injected at t=0 min. Projection of 3 Z-sections. **b**, Quantification of chromatin density before and after injection of AluI as in **a**. *n*=11 cells, 3 ROIs each. Significance tested by a paired, two-tailed *t*-test (*P*=0.013). **c**, Chromatin mobility in undigested metaphase chromosomes and after AluI injection, measured by fluorescence recovery after photobleaching in live metaphase cells expressing H2B-mCherry. Circles indicate photobleaching region at t=0 s. **d**, Quantification of fluorescence before and after photobleaching in *n*=8 (undigested) or *n*=10 (AluI-digested) cells as in **c**. **e**, AluI injection (t=0 min) during time-lapse microscopy as in **a**, but for a TSA-treated mitotic cell. **f**, Quantification of chromatin density as in **b,** normalized to the mean of untreated pre-injection cells shown in **b**. *n*=11 cells, 3 ROIs each. **g**, Chemical induction of G2-to-mitosis transition. HeLa cell expressing Aurora B-FRET biosensor was synchronized to G2 by RO3306 and then induced to enter mitosis by removing RO3306 and adding okadaic acid (OA). Mitotic entry was detected by chromosome compaction and FRET signal. Projection of 9 Z-sections. **h**, Quantification of chromatin density in G2 and mitosis as in **g** for *n*=13 cells. Bars indicate mean; significance was tested by a paired, two-tailed *t*-test (*P*=5.535×10^−12^). **i**, Chromatin was fragmented in G2 cells by injection of AluI and mitosis was subsequently induced as in **g**. Projection of 9 Z-sections. **j**, Quantification of chromatin density in G2 and mitosis as in **i** for *n*=8 cells. Bars indicate mean; significance was tested by a paired, two-tailed *t*-test (*P*=1.766×10^−7^). **k**, Chromatin was fragmented in TSA-treated G2 cells by injection of AluI and mitosis was subsequently induced as in **g**. Projection of 9 Z-sections. **l.** Quantification of chromatin density in G2 and mitosis as in **k** for *n*=10 cells. Bars indicate mean; significance was tested by a paired, two-tailed *t*-test (*P*=0.093). **m**, Microinjection of synthetic nucleosome arrays that were either untreated or pre-incubated with p300 acetyltransferase into live mitotic cells, for *n*=28 cells. Unmodified and acetylated nucleosome arrays were labelled by distinct fluorescent dyes. DNA was counterstained with DAPI. Bars indicate mean; significance was tested by a two-tailed Mann-Whitney test (*P*=1.645×10^−9^). Scale bars, **a**,**e**,**g**,**I**,**k**,**m**, 5 μm, insert 1 μm; **c**, 3 μm.

Intrinsic chromatin condensates dissolve following hyperacetylation^13^. To test whether acetylation also suppresses phase separation of mitotic chromosomes, we treated cells with TSA prior to mitotic entry and then microinjected AluI. This resulted in homogeneously dispersed chromatin fragments with almost no local condensates (Fig. 2e, f). In contrast, degradation of SMC4 had no effect on chromatin droplet formation after AluI-mediated fragmentation, indicating that condensins are not required to form an insoluble chromatin phase (Extended Data Fig. 4a, b). Thus, deacetylation is a major factor in establishing an immiscible chromatin phase in mitotic cells.

The observed differences in chromatin solubility might be due to variable efficiency of AluI-mediated DNA digestion in interphase or mitotic cells. To address this possibility, we used time-lapse microscopy to investigate whether chromatin fragments generated in interphase nuclei undergo a solubility phase transition upon progression to mitosis. To monitor cell cycle stages, we used a fluorescence resonance energy transfer (FRET) biosensor for a key mitotic kinase, Aurora B^37^. As chromosome fragmentation blocks mitotic entry owing to DNA damage signaling, we applied chemical inhibitors to induce an interphase-to-mitosis transition: we first synchronized cells to G2 using the Cdk1 inhibitor RO3306^38^ and then induced a mitosis-like state by removing RO3306 for Cdk1 activation and simultaneously inhibiting counteracting PP2A and PP1 phosphatases using Okadaic acid. Mitotic entry was demonstrated by the Aurora B FRET biosensor signal (Fig. 2g, h). Injection of AluI into G2 cell nuclei resulted in homogeneously distributed chromatin, consistent with a soluble state (Fig. 2h, i). Upon induction of mitosis, chromatin fragments formed spherical condensates that were as dense as intact chromosomes (Fig. 2h-j), whereas control cells in which RO3306 was not replaced by Okadaic acid maintained homogeneously dissolved chromatin fragments (Extended Data Fig. 4c d). These observations support a model in which a global reduction in chromatin solubility drives volume compaction at the interphase-to-mitosis transition.

To determine whether the loss of chromatin solubility upon mitotic entry depends on deacetylation, we inhibited histone deacetylases using TSA before injecting AluI into G2-synchronized cells. As in control cells, chromatin fragments homogeneously distributed throughout the nucleus in G2, but mitotic induction did not lead to the formation of condensed foci (Fig. 2k, l). Thus, active histone deacetylases are essential to form an immiscible chromatin phase upon mitotic entry.

Acetylation of chromatin might regulate solubility in cytoplasm directly or via other chromatin-associated components. To assess the effect of histone acetylation on chromatin solubility more specifically, we used synthetic nucleosome arrays^13^ as probes. We generated fluorescently labelled arrays of 12 unmodified naive nucleosomes as well as similar arrays that were labelled with a distinct fluorophore and acetylated *in vitro* with recombinant p 300 acetyltransferase. In vitro, the unmodified nucleosome arrays form liquid condensates under physiological salt concentrations, in contrast to the acetylated nucleosome arrays (Extended Data Fig. 4e, f)^13^. Following co-injection of these nucleosome arrays into live mitotic cells, unmodified arrays almost completely partitioned into the mitotic chromatin phase, whereas acetylated nucleosome arrays predominantly dissolved in the cytoplasm (Fig. 2m, n). Thus, histone acetylation is a direct regulator of chromatin solubility in cytoplasm. Overall, these results indicate that mitotic chromatin is a hydrogel in which condensin connects DNA loops and acetylation regulates chromatin solubility to mediate global compaction and formation of a sharp surface boundary.

### Chromatin condensates exclude negatively charged macromolecules

One explanation for how the immiscible mitotic chromatin phase acts as a barrier to microtubule penetration could be that it excludes cytoplasmic tubulin dimers and thereby limits microtubule growth at the cytoplasm-chromosome boundary. To investigate how soluble tubulin partitions relative to mitotic chromatin, we therefore microinjected fluorescently labelled tubulin into live mitotic cells and applied nocodazole to suppress microtubule polymerization (Fig. 3a). Soluble tubulin was much less concentrated inside mitotic chromosomes compared to the surrounding cytoplasm (Fig. 3a, b). In contrast, soluble tubulin was not excluded from hyperacetylated chromosomes in TSA-treated cells (Fig. 3a, b). Thus, the immiscible chromatin compartment formed by deacetylated mitotic chromatin excludes soluble tubulin.

**Figure 3.**
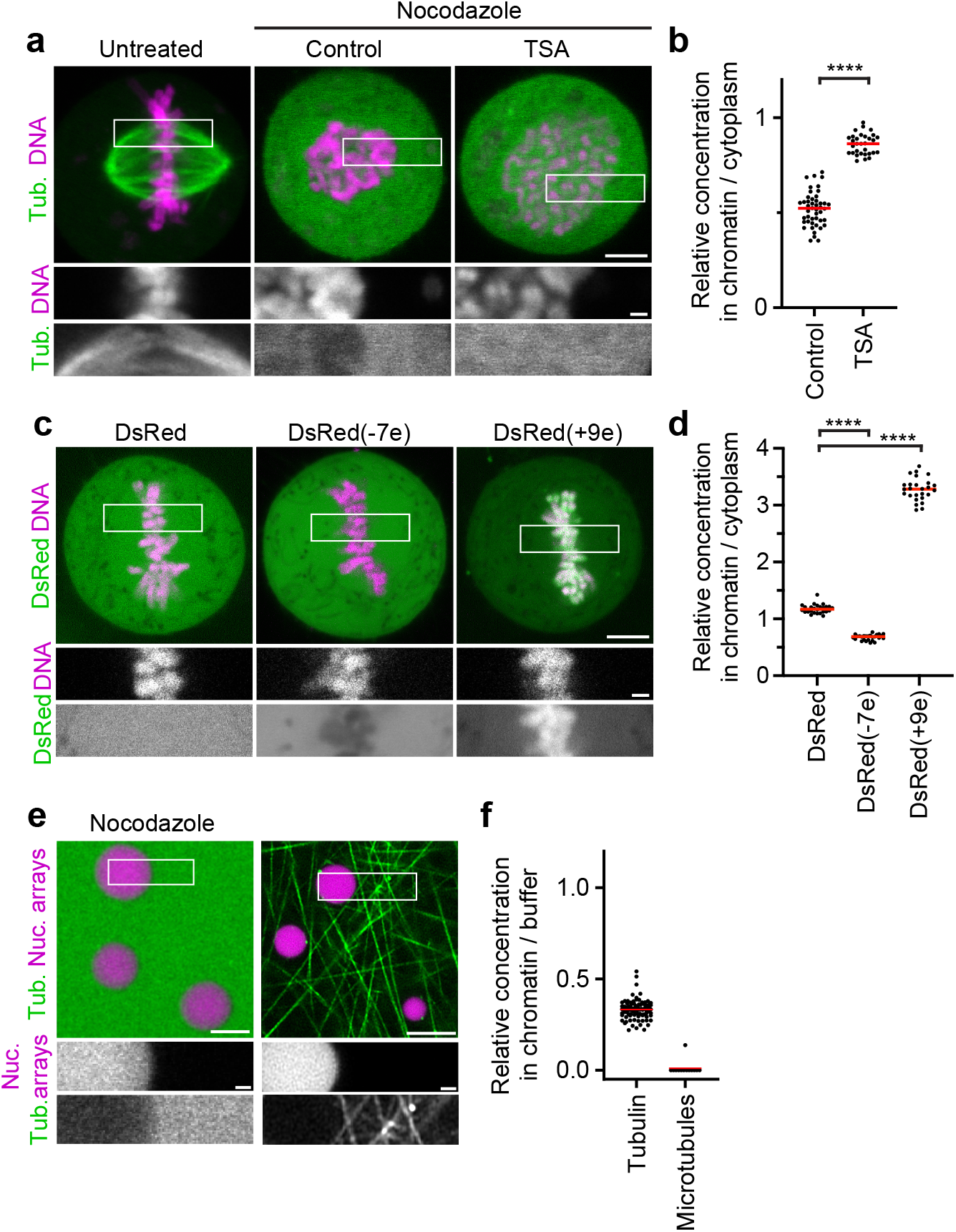
Chromatin condensates limit access of tubulin and other negatively charged macromolecules. **a**, Localization of tubulin relative to mitotic chromosomes. Rhodamine-labelled tubulin was injected into live mitotic cells that were either untreated, treated with nocodazole alone (Control) or in combination with TSA. **b**, Quantification of tubulin concentration for data shown in **a**. *n*=27 cells. Bars indicate mean; significance was tested by a two-tailed Welch *t*-test (*P*<1×10^−15^). **c**, Live cell images of HeLa cell expressing DsRed or DsRed fused at its N-terminus to electrically charged polypeptides. DNA stained with Hoechst. Numbers in brackets indicate predicted elementary charge of tetramers formed by DsRed fusion constructs. **d**, Quantification of DsRed concentration for data shown in **c**. *n*=26 cells (DsRed), *n*=26 (DsRed-7e), *n*=26 (DsRed+9e). Bars indicate mean; significances were tested by a two-tailed Welch *t*-test (DsRed(−7e), *P*<1×10^−15^; DsRed(+9e) *P*<1×10^−15^). **e**, Localization of tubulin relative to reconstituted nucleosome array droplets. Nucleosome array droplets were formed by incubation in phase separation buffer and fluorescently labelled tubulin was then added in the presence of nocodazole, or in the absence of nocodazole with subsequent temperature increase to 20 degrees to induce microtubule polymerization. **f**, Quantification of tubulin concentration or microtubule density in nucleosome array condensates relative to buffer for data shown in **e**. *n*=94 droplets, *n*=13 field of polymerized microtubules. Bars indicate mean. Scale bars, 5 μm, insert 1 μm.

To determine whether exclusion of tubulin is due to a limiting pore size in chromosomes, we expressed DsRed, a fluorescent protein that forms tetramers slightly larger than tubulin dimers^39,40^. In stark contrast to tubulin, DsRed distributed evenly across chromatin and cytoplasm (Fig. 3c, d). Thus, macromolecules in the size range of tubulin are not generally excluded from mitotic chromatin.

The selective exclusion of tubulin but not DsRed from mitotic chromatin suggests that specific molecular features control access. Tubulin is highly negatively charged at physiological pH (7.2), in contrast to near-neutrally charged DsRed, raising the possibility that macromolecular access is governed by electrostatic interactions. A high concentration of an overall negative electrical charge of chromatin^41^ in an immiscible condensate might therefore repel negatively charged cytoplasmic proteins. To investigate how electrical charge affects macromolecular access to mitotic chromatin, we fused charged polypeptides to the N-terminus of DsRed. DsRed fused to a negatively charged polypeptide (overall predicted charge on tetramer: −7e) was excluded from mitotic chromatin, whereas DsRed fused to a positively charged polypeptide (overall charge on tetramer: +9e) concentrated inside chromatin (Fig. 3c, d). Consistent with this observation, we found negatively charged mEGFP or dextrans were also excluded from mitotic chromosomes, whereas a positively charged surface mutant scGFP (super-charged GFP, +7e)^42^ or positively charged dextrans concentrated in mitotic chromosomes (Extended Data. Fig. 5a-d). Thus, electrical charge is a key determinant of macromolecular access to mitotic chromatin.

The exclusion of negatively charged macromolecules from mitotic chromosomes might be mediated by insoluble nucleosome fibers alone or it might involve other chromosome-associated factors. To investigate how tubulin interacts with pure chromatin condensates, we reconstituted droplets of recombinant nucleosome arrays *in vitro*^13^ and then added rhodamine-labelled tubulin in the presence of nocodazole. Soluble tubulin was indeed efficiently excluded from nucleosome array condensates (Fig. 3e, f). Consistent with this observation, negatively charged mEGFP or dextrans were also excluded from nucleosome array condensates, whereas a positively charged scGFP mutant or positively charged dextrans concentrated in nucleosome array condensates (Extended Data Fig. 5e-h). The exclusion of negatively charged macromolecules, including tubulin, is thus an intrinsic property of condensed nucleosome fibers.

The efficient exclusion of free tubulin suggests that weak affinity interactions in liquid chromatin condensates might be sufficient to limit microtubule polymerization. To investigate how microtubules interact with reconstituted chromatin droplets *in vitro*, we added purified tubulin and then induced microtubule polymerization by increasing the temperature. Microtubules formed a dense microtubule network, yet they almost never grew into chromatin condensates (Fig. 3e, f). Thus, chromatin-intrinsic material properties impose a highly impermeable barrier to microtubule polymerization independently of condensin or other chromosome-associated factors.

### Microtubules push liquid chromatin condensates

Microtubule polymerization exerts substantial pushing forces upon contact with stiff surfaces^43^. If the surface tension of the immiscible chromatin phase were strong enough to resist microtubule polymerization, then droplets of digested chromosomes should be pushed away by growing astral microtubules. To test this hypothesis, we injected AluI into mitotic cells treated with nocodazole and then washed out nocodazole to induce microtubule polymerization. We further applied S-trityl-L-cysteine (STLC)^44^ to maintain a monopolar astral spindle geometry. Time-lapse imaging showed that initially all chromatin resided in a single droplet at the cell center, but soon after nocodazole washout the chromatin split into several droplets that moved away from growing microtubule asters (Extended Data Fig. 6a, b). Thus, liquid chromatin condensates can be pushed by polymerizing astral microtubules.

Astral microtubules might directly push on chromatin droplets by polymerizing against the chromatin phase boundary or they might couple to chromatin via chromokinesins^19–21,45–47,23,22,48^. To investigate how the spindle moves chromatin droplets, we used RNAi to co-deplete two chromokinesins that are major contributors to the polar ejection force, Kid and Kif4a^45,47,23,22,48^ (Extended Data Fig. 6c-e). Following AluI-injection and nocodazole washout in the presence of STLC, chromosome droplets moved to the chromosome periphery as efficiently as in control cells (Extended Data Fig. 6c, d). Thus, spindle asters can move liquid chromatin independently of chromokinesins Kid and Kif4a, potentially by polymerizing microtubules pushing against the chromatin phase boundary.

Microtubule pushing on chromosome arms is counteracted by microtubule pulling at centromeres. To investigate how liquified chromatin responds to pulling forces at kinetochores, we injected AluI into cells expressing EGFP-tagged centromere-specific histone 3 variant CENP-A together with H2B-mCherry, in the presence of nocodazole. We then removed nocodazole to induce monopolar spindle assembly in the presence of STLC. The bulk mass of chromatin, visualized with H2B-mCherry, rapidly moved towards the cell periphery, while several much smaller chromatin condensates enriched in EGFP-CENP-A remained close to the spindle monopole (Fig. 4a, b). Thus, bulk chromatin is pushed by microtubules, while centromeres are transported towards the spindle pole. In cells after chromatin digestion, where the continuous connection between centromere and the remaining chromosome is lost, bulk chromatin and centromeric chromatin physically separate according to the locally dominating forces.

**Figure 4.**
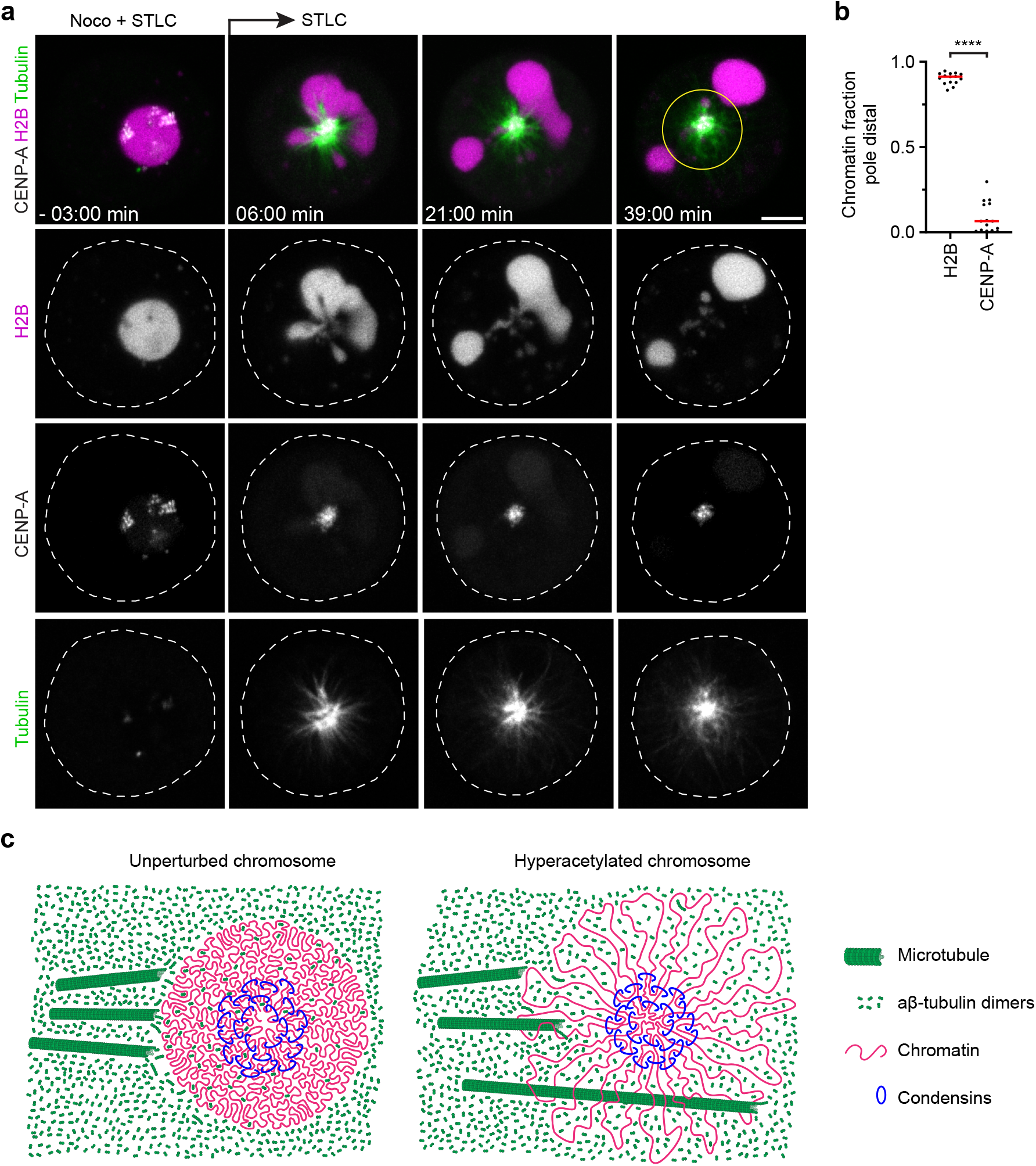
Microtubules push liquified chromatin away from the spindle pole. **a**, Time-lapse microscopy of liquified chromatin during monopolar spindle assembly. AluI was injected into live mitotic HeLa cells expressing H2B-mCherry and mEGFP-CENP-A, stained with SiR-tubulin, in the presence of nocodazole and STLC. Nocodazole was then removed at t=0 min during time-lapse imaging to induce monopolar spindle assembly. Projection of 5 Z-sections. **b**, Quantification of bulk chromatin (H2B-mCherry) and centromeric chromatin (mEGFP-CENP-A) localizing at the cell periphery relative to the region around the spindle monopole at t=36 min. *n*=15 cells. Bars indicate mean; significance was tested by a two-tailed Welch *t*-test (*P*<1×10^‑15^). **c**, Model of chromatin compaction and condensin-mediated DNA looping in mitotic chromosome and spindle assembly. Scale bars, 5 μm.

### Conclusions

Here, we show that a sharp global reduction in chromatin solubility converts chromosomes into phase-separated bodies during mitosis (Fig. 4c). The immiscible mitotic chromatin forms a surface dense in negative charge that prevents microtubule perforation and provides resistance to pushing forces while allowing chromatin fiber sliding internally, as required for continuous dynamic loop formation by condensin^6,14,49^. In parallel, condensin-mediated linkages establish a hydrogel that withstands pulling forces^14–16^. Jointly, these molecular activities shape discrete chromosome bodies with a defined surface despite continuous internal remodeling of the chromatin fiber.

Our results show that mitotic cells contain three principal domains with distinct microtubule polymerization propensity: the centrosome matrix, which concentrates soluble tubulin to promote nucleation at the poles^50^; the cytoplasm, which is highly permissive for microtubule growth and amplification; and a chromatin compartment that resists microtubule growth by exclusion of soluble tubulin and formation of a mechanical barrier towards polymerization. Our study thus provides a unified view of how chromatin looping by condensin and compaction through a volume phase transition contribute to the material properties and mechanical functions of mitotic chromosomes.

## Methods

### Cell lines and cell culture

All cell lines used in this study have been regularly tested negative for mycoplasm contamination. All cell lines in this study were derived from a HeLa “Kyoto” cell line previously described in ^51^. Cells were cultured in Dulbecco’s modified eagle medium (DMEM) (IMP/IMBA/GMI Molecular Biology Service/Media Kitchen) containing 10% (v/v) fetal bovine serum (FBS; Gibco, 10270-106, 2078432), 1% (v/v) penicillin-streptomycin (Sigma-Aldrich), 1% (v/v) GlutaMAX (Gibco; 35050038) and selected antibiotics according to the respective expression constructs: blasticidin S (6 μg mL^−1^, Thermo Fisher Scientific), puromycin (0.5 μg mL^−1^, Calbiochem), hygromycin B (0.25 mg mL^−1^) and G418 (1 mg mL-1, Invitrogen). HeLa cells were cultured at 37°C in a 5% CO2-containing atmosphere. Chromatin was visualized by stable expression of histone 2B labelled with mCherry (H2B-mCherry; Figs. 2a-f, 4, Extended Data Figs. 4a, b, 6a-d) or with Aurora B-FRET sensor (CFP/YFP) (Fig. 2g-l, Extended Data Fig. 4) or alternatively by labelling with Hoechst 33342 (1.62 μM, Invitrogen) (Figs. 1a-c, 2m, 3a-d, Extended Data Figs. 1a, 3c, 5a,c). Tubulin was visualized by stable expression of an N-terminally tagged eGFP-α-Tubulin fusion (Extended Data Figs. a, c) or by labeling with SiR-Tubulin (100 nM) (Figs. 1a, 4a, Extended Data Fig. 3c)^25^. Centromeres were visualized by stable expression of an N-terminally tagged eGFP-CENP-A fusion (Fig. 4a). Live-cell imaging was performed in DMEM containing 10% (v/v) FBS (Gibco, 10270-106, 2078432), 1% (v/v) penicillin-streptomycin (Sigma-Aldrich), and 1% (v/v) GlutaMAX (Gibco; 35050038) but omitting phenol red and riboflavin to reduce autofluorescence (imaging medium, short “IM”)^51^. Cells were grown in 10 cm or 15 cm CELLSTAR® (Greiner) dishes, in 75 cm^2^ or 175 cm^2^ Nunc™ EasYFlask™ cell culture flasks (Thermo Fisher Scientific) or in 96-well, 48-well, 12-well or 6-well Nunclon™ Delta Surface multi well plates (Thermo Fisher Scientific). For live-cell imaging, cells were cultivated on Nunc™ LabTek™ II chambered cover glass (Thermo Fisher Scientific), on μ-Slide 8 well covered coverslips (ibidi), in μ-Dish 35 mm, high, imaging dishes with polymer or glass bottom (ibidi).

### Generation of stable fluorescent reporter and genetically engineered cell lines

Cell lines stably expressing fluorescently labelled marker proteins were generated by random plasmid integration (for transfection conditions, see below) or by a lentiviral vector system pseudotyped with a mouse ecotropic envelope that is rodent-restricted (RIEP receptor system). Construction of RIEP receptor parental cell lines and subsequent generation of stable cell lines that express fluorescent marker proteins was performed as described previously^52^.

HeLa genome editing was performed using CRISPR-Cas9 mediated integration approach, using the chimeric Cas9-human Geminin fusion (Cas9-hGem) modification for enhanced editing efficiency^53^. Single-guide RNA (sgRNA) was cloned into pX330-U6-Chimeric_BB-CBh-hSpCas9-hGem-P2A-mCherry*aBpil (kind gift from Stefan Ameres). For tagging of endogenous Smc4 genes with an auxin-inducible degron tag, a repair template for targeting the C-terminus of the protein was designed, with homology flanks of 800 and 901 bp length for the 5’ and 3’ flanks, respectively. The repair template containing the coding sequence for a C-terminal mAID-HaloTag, including mutations of the protospacer adjacent motif (PAM), was synthesized as a gBlock® (IDT) and cloned into plasmid pCR2.1 for amplification. sgRNA/Cas9 and homology repair template plasmids were co-transfected using the Neon™ transfection system (Thermo Fisher Scientific). Transfection was performed according to the manufacturer’s guidelines (for HeLa cells: Pulse voltage, 1.005 V, pulse width, 35 ms, pulse number, 2), using the 100 μl tip cuvette, with 10 μg homology repair template and 10 μg gRNA/Cas9-hGEM for 1×10^6^ cells. 9 days after electroporation, cell pools were stained with Oregon Green® HaloTag (Promega, G2801) ligand and single clones isolated using fluorescence-activated cell sorting (FACS) and sorted into 96-well plates. Genotyping was performed as in a previous study^52^.

Integration of the OsTIR1(F74G) ligase into the adeno-associated virus integration site (AAVS1, designated “safe harbor”) locus for establishment of AID2 auxin-inducible degradation of Smc4 was achieved using a sgRNA and homology repair template strategy described in ^54^. Single-guide RNA was cloned into pX330-U6-Chimeric_BB-CBh-hSpCas9-hGem-P2A-mCherry*aBpil. For integration of the OsTIR1(F74G), a homology template containing an OsTIR1(F74G)-SnapTAG-MycA1-NLS and homology flanks of 804 and 837 bp length for the 5’ and 3’ flanks, respectively, was generated. Co-transfection of sgRNA/Cas9 and homology repair template plasmids was performed as described above. 8 days after electroporation, transfected cells were selected with 6μg/ml Blasticidin. Single clones were isolated from the stable pool by single cell dilution into 96-well plates. To identify clones which depleted SMC4-mAID-Halo, clones were treated for 90min with 1μM 5-Ph-IAA24, stained with Oregon Green® HaloTag (Promega, G2801) ligand and then analyzed by flow cytometry using an iQue Screener Plus.

### Transfection of plasmids and small interfering RNA

For expression of fluorescently labelled markers, the respective genes were cloned into bicistronic vectors containing an IRES or a 2A “self-cleaving” peptide sequence with antibiotic resistance genes that allow the protein of interest and the resistance gene to be expressed from the same transcript. For transient transfection or transfection for subsequent selection and colony picking, plasmids were transfected using X-tremeGENE 9 DNA transfection reagent (Roche, 6365787001) following the manufacturer’s protocol (1 μg plasmid, 4.5 μl X-tremeGENE 9 in 100 μl serum-free OptiMEM) or PEI transfection reagent (1 mg ml^−1^ stock, Polyscience 24765-2, 4 μg of transfection reagent per 1 μg of plasmid). For stable expression, plasmids were transfected using PEI, 10 μg plasmid for one ~70 confluent 15 cm dish and incubated for 48 h before antibiotic selection.

Small interfering RNAs (siRNAs) were delivered with lipofectamine RNAiMax (Invitrogen) according to the manufacturer’s instructions. hKid (Kif22) was targeted using 16 nM custom silencer select siRNA (sense strand CAAGCUCACUCGCCUAUUGtt, Thermo Fisher Scientific, including a 3’ overhang tt dinucleotide for increased efficiency). Kif4A was targeted using 16 nM custom silencer s elect siRNA (sense strand GCAAGAUCCUGAAAGAGAUtt, Thermo Fisher Scientific, including a 3’ overhang tt dinucleotide for increased efficiency). Custom silencer select siXWNeg (sense strand UACGACCGGUCUAUCGUAGtt, Thermo Fisher Scientific, including a 3′ overhanging tt dinucleotide for increased efficiency) was used as a non-targeting siRNA control. Cells were imaged 30 h after siRNA transfection. Co-transfection of hKid and Kif4a siRNAs was performed using 16 nM final concentration each. Both hKid and Kif4a siRNAs were described in ^23^.

### Inhibitors and small molecules

To degrade Smc4-mAID-Halotag, cells were incubated in 1 μM 5-Phenylindole-3-acetic acid (5-PhIAA) (Bio Academia, 30-003) for 2.5-3h. To induce histone hyperacetylation, cells were incubated with 5 μM Trichostatin A (TSA) (Sigma-Aldrich, T8552). To arrest cells in prometaphase with monopolar spindle configuration, cells were incubated in 5 μM S-Trityl-L-cysteine (STLC) (Enzo Life Sciences, ALX-105-011-M500) for 2-3h. To arrest cells in prometaphase, cells were incubated for 2-3h in nocodazole (200 ng ml^−1^ for live-cell imaging only)or 100 ng ml^−1^ for microinjection and/or subsequent washout; Sigma-Aldrich, M1404). Nocodazole washout was by washing cells 5 times with prewarmed imaging medium on the microscope. To arrest cells in metaphase for microinjection, cells were incubated with 12 μM proTAME (R&D systems, I-440-01M) for 2 h. To synchronize cells to the G2-M boundary, cells were first synchronized with a double thymidine block followed by a single RO3306 block. One day after seeding, cells were transferred into medium containing 2 mM thymidine (Sigma-Aldrich, T1895). 16 h later, cells were washed three times with prewarmed medium and released for 8 h. The thymidine block was repeated once. 6 h after the second thymidine release cells were transferred into medium containing 8 μM RO3306 (Sigma-Aldrich, SML0569) for 4 h. Following G2 arrest, RO3306 was removed and cells were washed three times with imaging medium containing 1.5 μM okadaic acid (LC laboratories, O-5857).

### Immunofluorescence

For all immunofluorescence experiments, cells were grown on sterilized 18 mm round Menzel cover glasses in 24-well multi-well plates, except immunofluorescence against acetylated histones, which was performed on Nunc™ LabTek™ II chambered cover glass. For co-staining spindle poles and centromeres, cells were fixed and extracted at the same time with 1XPHEM (60 mM K-PIPES (Sigma-Aldrich), pH 6.9, 25 mM K-HEPES (Sigma-Aldrich), pH 7.4, 10 mM EGTA (Merck, 324626), 4 mM MgSO_4_•7H_2_O (Sigma-Aldrich)) containing 0.5% Tween-20 (Sigma-Aldrich) and 4% methanol-free formaldehyde (Thermo Fisher Scientific) for 10 min. The fixation reaction was quenched with 10 mM Tris-HCl (Sigma-Aldrich), pH 7.4, in phosphate-buffered saline (PBS), washed again with PBS and blocked in 10% normal goat serum (Abcam), 0.1%Tween-20 (Sigma-Aldrich) for 1 h.

For staining of acetylated histone tails or CyclinB1, cells were fixed in PBS containing 4% methanol-free formaldehyde (Thermo Fisher Scientific, 28906) for 10 min. The fixation was quenched with 10 mM Tris-HCl (Sigma-Aldrich) pH 7.4 in PBS for 5 min, and cells permeabilized with PBS containing 0.5% Triton-X100 for 15 min, washed again with PBS and blocked using 2% bovine serum albumin (BSA) (Sigma-Aldrich, A7030) in PBS, containing 0.2% Tween-20 (Sigma-Aldrich), for 1 h. Antibody dilution buffer was composed of PBS with 2% BSA (Sigma-Aldrich, A7030), containing 0.1% Tween-20 (Sigma-Aldrich). After primary and secondary antibody incubations, samples were washed with PBS containing 0.1% Tween-20 (Sigma-Aldrich), three times for 10 min each time. CENP-A was detected with a monoclonal mouse antibody (Enzo Life Sciences, ADI-KAM-CC006-E, 10161910, 1:1000) and visualized using a goat anti-mouse Alexa Fluor 488 secondary antibody (Molecular Probes, A11001, 1:1000) (Extended Data Fig. 1d). Pericentrin was detected with a recombinant rabbit antibody (Abcam, ab220784, GR3284309-1, 1:2000) and visualized using a goat anti-rabbit Alexa Fluor 633 secondary antibody (Molecular Probes, A21071, 1:1000) (Extended Data Fig. 1d). Acetylated histone2B was detected using a polyclonal rabbit antibody (Millipore, 07-373, 3092508, 1:500) and visualized using a donkey anti-rabbit Alexa Fluor 488 secondary antibody (Molecular Probes, A21206, 1:1000) (Extended Data Figs. 2a, c, 3a). Acetylated histone 3 was detected using a polyclonal rabbit antibody (Merck, 06-599, 3260200, 1:500) and visualized using a donkey anti-rabbit Alexa Fluor 488 secondary antibody (Molecular Probes, A21206, 1:1000) (Extended Data Fig. 2a, c). Acetylated Histone 4K16 was detected using a recombinant rabbit antibody (Abcam, ab177191, GR284778-8, 1:400) and visualized using a donkey anti-rabbit Alexa Fluor 488 secondary antibody (Molecular Probes, A21206, 1:1000) (Extended Data Figs. 2a, c). Cyclin B1 was detected with a monoclonal rabbit antibody (Cell Signaling, 12231S, 7, 1:800) and visualized using a donkey anti-rabbit Alexa Fluor 488 secondary antibody (Molecular Probes, A21206, 1:1000) (Extended Data Fig. 2e). Immunofluorescence samples prepared in Nunc™ LabTek™ II chambered cover glass wells were stored in PBS containing 1.62 μM Hoechst 33342 (Invitrogen). Immunofluorescence samples prepared on 18 mm round Menzel cover glasses were embedded in Vectashield with or without DAPI (Vectorlabs, H-1000 or H-1200).

### Preparation of 384-well microscopy plates and coverslips for *in vitro* assays

384-well μCLEAR® microscopy plates (Greiner Bio-One, 781906) were washed with 5% HellmanexIII (Lactan, 105513203) in ≥18 MΩ MonoQ H_2_O at 65°C in a table top Incu-Line oven (VWR) for 4 h and then rinsed 10 times with ≥ 18 MΩ MonoQ H_2_O. Silica was etched with 1 M KOH (Sigma-Aldrich) for 1 h at room temperature rinsed again 10 times with ≥ 18 MΩ MonoQ H_2_O. Etched multiwell plate was treated with 5k-mPEG-silane (Creative PEGworks, PLS-2011) suspended in 95% ethanol (VWR) for 18h at room temperature. Plate was washed once with 95% EtOH, then 10 times with ≥ 18 MΩ MonoQ H_2_O and dried in a clean chemical hood overnight. Until individual wells were used, the plate was sealed with adhesive PCR plate foil (Thermo Fisher Scientific) and kept in a dry and dark space. Before an experiment, foil above individual wells was cut and 50 μl of 100 mg ml^−1^ BSA (Sigma-Aldrich, A7030).

For microtubule/nucleosome array droplet experiments (Fig. 3e), thin layer cover glass sandwiches were constructed from passivated 24×60 mm Menzel coverglass (VWR). To clean coverslips, they were vertically stacked into a Coplin jar (Canfortlab, LG084). Coplin jars were then sonicated in acetone (Sigma-Aldrich) for 15 min, in 100% EtOH (VWR) for 15 min and then washed 10 times with ≥ 18 MΩ MonoQ H_2_O. All sonication steps were performed using an ultrasonic cleaning bath (Branson, 2800 Series Ultrasonic Cleaner, M2800). In a separate Erlenmeyer flask, 31.5 ml 30% aqueous hydrogen peroxide solution (Sigma-Aldrich, 31642) was added to 58.5 ml concentrated sulfuric acid (Sigma-Aldrich, 258105) (“piranha” acid). The flask was gently swirled until bubbling and heat occurred, then the whole contents of the flask was added to the coverslips in the Coplin jars, ensuring that all coverslip surfaces were covered. The jar was transferred into a 95°C water bath and heated for 1 h. Afterwards, piranha solution was discarded, and cover glasses washed once with ≥18 MΩ MonoQ H_2_O and etched with 0.1 M KOH for 5 min. Cover glasses were transferred to a fresh, dry Coplin jar and dried to completion in a 65°C bench top oven, and afterwards let to cool to room temperature. In a separate Erlenmeyer flask, 4 ml dichlorodimethylsilane (DCDMS) (Sigma-Aldrich, 440272, anhydrous) was injected into 80 ml heptane (Sigma, 246654, anhydrous). The contents of the flask were immediately transferred to the jar containing the cover glasses, and jar was incubated for 1 h at room temperature. Afterwards, silane was decanted, and cover glasses are sonicated in chloroform (Sigma-Aldrich) for 5 min, washed in chloroform once, sonicated in chloroform again for 5 min, and sonicated twice in ≥18 MΩ MonoQ H_2_O. Finally, cover glasses were washed once more in chloroform, air dried, and stored in a sealed container in a dry and dark, dust-free space (up to 6 months).

To passivate cover glasses, on the day of an experiment a cover glass was transferred to a drop of 5% Pluronic F-127 (Thermo Fisher Scientific, P6866) dissolved in BRB80 buffer (80 mM K-PIPES, pH 6.9 (Sigma-Aldrich, P6757), 1 mM MgCl_2_ (Sigma-Aldrich), 1 mM EGTA (Merck, 324626)) for >2 h at room temperature. Directly prior to assembly of the imaging chamber, cover glass was rinsed once with ≥18 MΩ MonoQ H_2_O and once with BRB80 (80 mM K-PIPES, pH 6.9 (Sigma-Aldrich, P6757), 1 mM MgCl_2_ (Sigma-Aldrich), 1 mM EGTA (Merck, 324626)).

### Expression and purification of GFP proteins

pET-based constructs for the expression and purification of mEGFP(−7e) and scGFP(+7e) were a generous gift from Dr. David Liu^55^. An overnight culture of Rosetta 2 (pLysS) *E. coli* (Novagen) transformed with pET_-7GFP, or pET_+7GFP plasmids encoding GFP with a theoretical peptide charge of −7e or +7e, respectively, were grown on an agar plate by re-plating a single transformant on LB supplemented with 100 ng/μl of ampicillin at 37°C. The bacterial lawn was suspended in LB supplemented with the 100 ng/μl of ampicillin and grown at 37°C to a density (OD_600_ nm) of 0.4, cooled over 1 h to 18°C, and recombinant protein expression was induced by addition of IPTG to 0.5 mM for 18 h at 18°C. The cells were collected by centrifugation, resuspended in NiNTA Lysis Buffer (50 mM HEPES•NaOH, pH 7, 150 mM NaCl, 10% [w/v] glycerol, 15 mM Imidazole, 5 mM ß-mercaptoethanol, 1 mM Benzamidine, 100 μM Leupeptin, 100 μM Antipain, 1 μM Pepstatin), the cellular suspension was flash frozen in liquid N_2_, and stored at −80°C.

Bacterial cultures with expressed GFP proteins in NiNTA Lysis Buffer were thawed in a water bath and lysed by multiple passages through an Avestin Emulsiflex-C5 high pressure homogenizer at ~10,000 PSI. An equal volume of NiNTA Dilution Buffer (50 mM HEPES•NaOH, pH 7, 1.85 M NaCl, 10% [w/v] glycerol, 15 mM Imidazole, 5 mM ß-mercaptoethanol, 1 mM Benzamidine, 100 μM Leupeptin, 100 μM Antipain, 1 μM Pepstatin) was added to the lysate to increase NaCl concentration to 1 M. Soluble bacterial lysate was isolated by centrifugation of cellular debris in a Beckman Avanti J-26 XPI centrifuge in a JA25.5 rotor at 20,000 RPM. Soluble lysate was incubated with NiNTA resin (Qiagen) equilibrated in NiNTA Wash Buffer A (50 mM HEPES•NaOH, pH 7, 1 M NaCl, 10% [w/v] glycerol, 15 mM Imidazole, 5 mM ß-mercaptoethanol, 1 mM Benzamidine, 100 μM Leupeptin, 100 μM Antipain, 1 μM Pepstatin) for 2 h in batch with end-over-end mixing. NiNTA resin was poured into a BioRad EconoColumn and resin was washed 20 column volumes of NiNTA Wash Buffer A and 20 column volumes of NiNTA Wash Buffer B (50 mM HEPES•NaOH, pH 7, 150 mM NaCl, 10% [w/v] glycerol, 15 mM Imidazole, 5 mM ß-mercaptoethanol, 1 mM Benzamidine, 100 μM Leupeptin, 100 μM Antipain, 1 μM Pepstatin) before elution in NiNTA Elution Buffer (50 mM HEPES•NaOH, pH 7, 150 mM NaCl, 10% [w/v] glycerol, 350 mM Imidazole, 5 mM ß-mercaptoethanol, 1 mM Benzamidine, 100 μM Leupeptin, 100 μM Antipain, 1 μM Pepstatin).

GFP proteins were diluted with 9 volumes of Buffer SA (20 mM HEPES•NaOH, pH 7, 10% [w/v] glycerol, 1 mM DTT) and applied to Source 15S (meGFP) or Source15Q (scGFP) resin (GE Healthcare) equilibrated in 98.5% Buffer SA and 1.5 % Buffer SB (20 mM HEPES•NaOH, pH 7, 1 M NaCl, 10% [w/v] glycerol, 1 mM DTT) and eluted with a linear gradient to 100% Buffer SB. Fractions containing GFP proteins were concentrated in a centrifugal concentrator with a 3,000 dalton MWCO and purified further by s i ze exclusion chromatography using a Superdex 200 10/300 GL gel filtration column equilibrated with Gel Filtration Buffer (20 mM Tris•HCl, pH 8, 150 mM NaCl, 10% [w/v] glycerol, 1 mM DTT). Peak fractions of purified mEGFP(−7e) and scGFP(+7e) protein were concentrated in a centrifugal concentrator, as above, and quantified by measuring protein absorbance at 280 nm using their calculated molar extinction coefficient (https://web.expasy.org/protparam/) of 23,380/ M•cm for meGFP and 18,910/M•cm for scGFP. Purified proteins were flash frozen in liquid N2 and stored at −80°C in single use aliquots.

### Nucleosome array *in vitro* experiments

Generation, fluorophore labelling, assembly and quality control of nucleosome arrays has been described in ^13^. In this study, 12X601 nucleosome arrays carrying no label, an Alexa Fluor 488, or Alexa Fluor 594 label were used. The covalently fluorescently labelled nucleosome arrays used for microinjection were labeled at an 8% fluorophore density (8 in 100 histone2B proteins labeled with a fluorophore) and dialyzed against TE (10 mM Tris-HCl (Sigma-Aldrich), pH 7.4 (Sigma-Aldrich), 1 mM EDTA (Sigma-Aldrich), pH 8.0, 1mM DTT (Roche, 10708984001)) to remove glycerol.

To induce phase separation of nucleosome arrays (Extended Data Figs. 4e, 5e,g), arrays were first equilibrated in chromatin dilution buffer (25 mM Tris•OAc, pH 7.5 (Sigma-Aldrich), 5 mM DTT (Roche, 10708984001), 0.1 mM EDTA (Sigma-Aldrich), 0.1 mg ml^−1^ BSA (Sigma-Aldrich, A7030), 5% (w/v) glycerol (Applichem, A0970)) in the presence of 2.03 μM Hoechst 33342 (Invitrogen, H1399) when using unlabeled arrays for 10 min at room temperature. Phase separation was induced by addition of 1 volume phase separation buffer (25 mM Tris•OAc, pH 7.5 (Sigma-Aldrich), 5 mM DTT (Roche, 10708984001), 0.1 mM EDTA (Sigma-Aldrich), 0.1 mg ml-1 BSA (Sigma-Aldrich, A7030), 5% (w/v) glycerol (Applichem, A0970), 300 mM KOAc (Sigma-Aldrich), 1 mM Mg[OAc]2, 2 μg ml^−1^ glucose oxidase (Sigma-Aldrich, G2133), 350 ng ml^−1^ catalase (Sigma-Aldrich, C1345) and 4 mM glucose (AMRESCO, 0188)). For in vitro assays containing tubulin, EGTA was substituted for EDTA in the dilution and phase separation buffers. After addition of phase separation buffer, the final concentration of nucleosome arrays per reaction was 500 nM, and reactions were incubated at room temperature for 10 min before transferring the suspension to imaging chambers.

To visualize soluble tubulin partitioning relative to chromatin droplets *in vitro*, 20 μl of chromatin droplet suspension were transferred to a passivated well of a 384-well microscopy plate. Chromatin droplets were to sedimented for 15 min. Afterwards, 5 μl of soluble TRITC-labelled tubulin (Cytoskeleton, TL590M) in phase separation buffer containing 500 ng ml^−1^ nocodazole was added for a final tubulin concentration of 5 μM. Soluble tubulin equilibrated for 10 min before images were recorded.

To visualize GFP surface charge variant partitioning relative to chromatin droplets *in vitro*, 20 μl of chromatin droplet suspension were transferred to a passivated well of a 384-well microscopy plate. Chromatin droplets were allowed to sediment for 15 min, after which 5 μl of GFP solution in phase separation was added to a final concentration of 5 μM. GFP proteins equilibrated for 10 min before images were recorded.

To visualize chemically modified dextran partitioning into chromatin droplets *in vitro*, 20 μl of chromatin droplet suspension were transferred to a passivated well of a 384-well microscopy plate. Chromatin droplets were allowed to sediment for 15 min, after which 5 μl dextran solution in phase separation buffer was added to a final concentration of 500 μg ml^−1^. The dextran used was a fluorescein (FITC) labelled 4.4 kDa dextran fraction (negative overall charge conferred to dextran by fluorophore charge) (Sigma-Alrich, FD4) or a FITC labelled 4.4 kDa dextran fraction modified with diethylaminoethyl (DEAE) groups conferring overall positive charge (TdB, DD4).

All images of macromolecule partitioning relative to chromatin droplets *in vitro* (Fig. 3e, Extended data Fig. 5e, g) were recorded on the incubated stage of a customized Zeiss LSM980 microscope using a x40 1.4 NA Oil DIC Plan-Apochromat objectives (Zeiss).

### Nucleosome array acetylation *in vitro*

To generate acetylated nucleosome arrays for microinjection, recombinant p300 histone acetyl transferase domain was generated (VBCF Protein technologies Facility) using plasmid pETduet+p300_HAT_ according to a purification strategy described in ^13^. To induce histone acetylation of nucleosome arrays, arrays with 8% Alexa Fluor 488 or Alexa Fluor 594 label density at a concentration of 3.85 μM were incubated with 6.12 μM p300-HAT (enzyme stock 61.2 μM in gel filtration buffer (20 mM Tris•HCl, pH 8.0, 150 mM NaCl, 10% (w/v) glycerol, 1mM DTT)) in the presence of 750 μM Acetyl-CoA (Sigma-Aldrich, A2056) in gel filtration buffer at room temperature for 2 h with occasional agitation. Afterwards, the acetylation was stopped by addition of A-485 (Tocris, 6387) to a final concentration of 9 μM and the reaction mixture stored in the dark. To 10 μl of quenched acetylation reaction, 10 μl dilution buffer containing 5 μM A-485 (Tocris, 6387) were added and reaction allowed to equilibrate at room temperature for 10 min. Afterwards, 1 volume of phase separation buffer containing 5 μM A-485 (Tocris, 6387) was added and the mixture incubated for 10 min at room temperature before transferring the suspension to a passivated 384-well microscopy plate well.

### Microtubule *in vitro* polymerization

To generate stabilized microtubule seeds for nucleation of microtubule networks *in vitro*, 22 μl of 5 mg ml^−1^ tubulin protein (Cytoskeleton, T240) was mixed with 2 μl of 5 mg ml^−1^ TRITC-labeled tubulin protein (Cytoskeleton, TL590M) and 1 μl of 5 mg ml^−1^ Biotin-XX-labeled tubulin (Pursolutions, 033305). All tubulin storage solutions were prepared in BRB80 (80 mM K-PIPES, pH 6.9 (Sigma-Aldrich, P6757), 1 mM MgCl_2_ (Sigma-Aldrich), 1 mM EGTA (Merck, 324626)). The tubulin mixture was centrifuged at 4°C for 15 min in a tabletop centrifuge (Eppendorf, 5424R) at 21k x g. To 22.5 μl of the resulting supernatant, 2.5 μl 10 mM guanylyl-(alpha, beta)-methylene-diphosphonate (GMPCPP) (Jena Biosciences, NU-405) were added to a final concentration of 1 mM and the resulting solution incubated at 37°C in a water bath in the dark for 30 min. The resulting seeds were sheared using a 22-gauge needle Hamilton syringe (Sigma-Aldrich, 20788). The resulting suspension was stored in the dark. Seeds were prepared freshly per day and used for subsequent experiments.

Soluble tubulin polymerization mix containing 50 μM tubulin protein (Cytoskeleton, T240), 1 μM TRITC-labeled tubulin protein (Cytoskeleton, TL590M) and 1 mM GTP (Sigma-Aldrich, G8877) in BRB80 (80 mM K-PIPES, pH 6.9 (Sigma-Aldrich, P6757), 1 mM MgCl_2_ (Sigma-Aldrich), 1 mM EGTA (Merck, 324626)) was centrifuged at 4°C for 15 min in a tabletop centrifuge (Eppendorf, 5424R) at 21k x g. The resulting supernatant was used to assemble a microtubule polymerization mix containing 20 μM total soluble tubulin (19.61 μM unlabeled tubulin dimers, 0.39 μM labelled tubulin), 50 mM DTT (Roche, 10708984001), 120 μM glucose (AMRESCO, 0188), 1 mM GTP (Sigma-Aldrich, G8877), 20 μg ml^−1^ glucose oxidase (Sigma-Aldrich, G2133), 175 ng ml^−1^ (Sigma-Aldrich, C1345) in BRB80 (80 mM K-PIPES, pH 6.9 (Sigma-Aldrich, P6757), 1 mM MgCl_2_ (Sigma-Aldrich), 1 mM EGTA (Merck, 324626)). Composition of soluble tubulin and tubulin polymerization mix was adapted from a procedure for visualization of microtubule plus-end tracking (+TIP) proteins^56^.

To visualize chromatin droplets along with polymerized microtubules, imaging chambers were constructed as described in ^57^. Prior to Pluronic F-127 passivation, each silanized 24×60 mm cover glass was cut into a 24×25 mm (top piece) and 24×35 mm (bottom piece) using a diamond tipped steel scribe (Miller, DS-60-C). The bottom coverslip was mounted into the support slide and fixed in place with preheated VALAP (1 part Vaseline (Sigma-Aldrich, 16415), 1 part lanolin (Sigma-Aldrich, L7387), 1 part Paraffin wax (Sigma-Aldrich, 327204); all parts by weight). In the central region of the bottom cover glass, a 2 Well Silicon Culture-Insert (ibidi, 80209) was attached and 40 μl of nucleosome array droplet suspension added to one of the wells. The suspension was sedimented for 15 min in a humidified chamber at room temperature. Afterwards, 32 μl of buffer were removed from the well, the Culture-Insert removed from the coverslip and 20 μl soluble tubulin mix added to the remaining chromatin droplet suspension. The top coverslip was added on top with the passivated side facing the reaction mixture, and the droplet was allowed to spread fully (~10 s). Afterwards, the sample was sealed with VALAP to prevent evaporation and thermal streams within the liquid film. This procedure yielded a liquid film of ~30 μm thickness. The assembled imaging cell was transferred to the incubated stage of a customized Zeiss LSM980 microscope combined with the Airyscan2 detector, using a x63 1.4 NA Oil DIC Plan-Apochromat objective (Zeiss). The sample was incubated at 37°C for >30 min before images were recorded.

### CLEM and electron tomography

HeLa cells stably expressing H2B-mCherry were cultured on carbon-coated Sapphire discs (0.05 mm thick, 3 mm diameter; Wohlwend GmbH). To enrich prometaphase cells, cells were synchronized to G2-M using a double thymidine (Sigma-Aldrich) block for 16 h with 2 mM thymidine each and a subsequent RO3306 (Sigma-Aldrich) block for 6 h with 8 μM RO3306. During the RO-3306 block, cells were treated with 5 μM TSA for 3 h prior to RO3306 washout. RO3306 was washed out by rinsing the cells 3 times with prewarmed imaging medium. Cells were observed on a customized Zeiss LSM780 microscope, using a x20 0.5 NA EC PlnN DICII Air objective (Zeiss). 25 min after the RO3306 washout, most cells reached prometaphase (based on DIC and chromatin morphology), and cells were subsequently processed for electron microscopy. Immediately before freezing, cells were immersed in cryoprotectant solution (imaging medium containing 20% Ficoll PM400; Sigma Aldrich), and instantly frozen using a high-pressure freezing machine (HPF Compact 01; Wohlwend GmbH).

Frozen cells were freeze-substituted into Lowicryl HM20 resin (Polysciences Inc., 15924-1) using freeze-substitution device (Leica EM AFS2, Leica Microsystems) as follows: Cells were incubated with 0.1% uranyl acetate (UA) (Serva Electrophoresis GmbH, 77870) in acetone at −90 °C for 24 h. The temperature was increased to −45 °C at a rate of 5 °C/h and then incubate for 5 h at −45 °C. Cells were washed three times in acetone at −45°C and then incubated in increasing concentrations of resin Lowicryl, HM20 in acetone (10, 25, 50, and 75% for 2 h each) while the temperature is further increased to −25°C at a rate of 2.5°C/h. Cells were then incubated in 100% resin Lowicryl, HM20 at −25°C for a total of 16 h, changing the pure resin solution after 12 h, 14 h, and 16 h. The resin was polymerized under UV light at −25 °C for 48 h. The temperature was increased to 20°C (5°C/h) and UV polymerization was continued for another 48 h.

After resin polymerization, sections of 250 nm thickness were cut with an ultramicrotome (Ultracut UCT; Leica) and collected on copper–palladium slot grids (Science Services) coated with 1% Formvar (Plano GmbH). The sections were post-stained with 2% UA in 70% methanol at room temperature for 7 min and lead citrate at room temperature for 5 min. The sections were observed by Tecnai F20 transmission EM (200 kV; FEI). H2B fluorescence was used to choose cells and subcellular ROIs for tomography. For tomography, a series of tilt images were acquired over a −60° to +60° tilt range with an angular increment of 1° using Serial EM software^58^ at a final x-y pixel size of 1.14 nm. Tomograms were reconstructed using R-weighted back projection method implemented in the IMOD software package^59^.

### Western blotting

Samples were separated by Novex NuPAGE SDS-PAGE system (Thermo Fisher Scientific) using 4% - 12% BisTris in MES running buffer according to manufacturer’s guidelines, and transferred to a 0.45 μm nitrocellulose membrane (Bio-Rad) by tank-blot wet transfer (Bio-Rad) at room temperature for 1 h. Blocking, primary antibody incubations (4°C, 16 h) and secondary antibody incubations (room temperature, 1.5 h) were performed in 5% (w/v) milk (Maresi, “fixmilch” Instant milk powder) in PBS, containing 0.05% Tween-20 (Sigma-Aldrich). Smc4 was probed using a rabbit polyclonal antibody (Abcam, ab229213, GR3228108-5, 1:1000). GAPDH was probed with a rabbit polyclonal antibody (Abcam, ab9485, GR3212164-2, 1:2500). hKid was probed with a monoclonal rabbit antibody (Abcam, ab75783, GR129278-4). Kif4a was probed with a recombinant rabbit antibody (Abcam, ab124903, GR96215-7, 1:1000). Horseradish peroxidase (HRP)-conjugated anti-mouse or anti-rabbit secondary antibodies (Bio-Rad, 1:10000) were visualized using ECL Plus Western Blotting Substrate (Thermo Fisher Scientific) on a Bio-Rad ChemiDoc Imager.

### Microscopy

Images of G2-to-mitosis inductions, DsRed transfections, HaloTag stainings of wildtype and Smc4-mAID-HaloTag cells, histone acetylation, p300 overexpression, and fluorescence recovery after photobleaching (FRAP) experiments were recorded on a customized Zeiss LSM780 microscope, using x40 or x63 1.4 NA Oil DIC Plan-Apochromat objectives (Zeiss), operated by ZEN Black 2011 software. Images of chromatin density, live-microtubule stains, AluI-digestion time-lapse videos, nucleosome array microinjections, tubulin microinjections, CENP-A/Pericentrin immunofluorescence experiments and Cyclin B1 staining were recorded on a customized Zeiss LSM980 microscope combined with the Airyscan2 detector, using x40 or x63 1.4 NA Oil DIC Plan-Apochromat objectives (Zeiss), operated by ZEN3.3 Blue 2020 software. For all confocal microscopes, an incubator chamber (EMBL) provided a humidified atmosphere and a constant 37 °C temperature with 5% CO2.

For FRAP experiments, selected image regions were bleached using a laser intensity 200-fold higher than the laser intensity used for image acquisition, and each bleached pixel was illuminated 20 times with the pixel dwell time used for acquisition (1.79 μs). Images were acquired every 25 ms for the course of the experiment.

Images of nucleosome array droplets and *in vitro* polymerized microtubules were recorded on a customized Zeiss LSM980 microscope combined with the Airyscan2 detector, using x40 or x63 1.4 NA Oil DIC Plan-Apochromat objectives (Zeiss), operated by ZEN3.3 Blue 2020 software. For mounting 384-well microscopy plates, a Pecon® Universal Mounting Frame KM adapter was used. For mounting microtubule imaging slides, a custom aluminum mounting block for 24 mm X 24-60 mm coverslips was used (IMP/IMBA workshop and BioOptics).

### Microinjection experiments

Live-cell microinjection experiments were performed using a FemtoJet® 4i microinjector in conjunction with an InjectMan® 4 micromanipulation device (Eppendorf). All microinjections were performed using pre-pulled Femtotips® injection capillaries (Eppendorf). The microinjection device was directly mounted on a customized confocal Zeiss LSM780 (Figs. 2c, g,i,l, Extended Data Fig. 4c) or a customized Zeiss LSM980 microscope (Figs. 2a, e, m, 3a, 4a, Extended Data Figs. 4a, 5a, c, 6a, c) with live-cell incubation (37°C, 5% CO2). For all microinjections, cells were cultured in μ-Dish 35 mm high wall imaging dishes with polymer or glass bottom (ibidi) to reach near-confluency on the day of the injection.

For injection of AluI (Fast Digest, Thermo Fisher Scientific, FD0014), 1 volume of AluI-stock was added to 2 volumes of 5 mg ml^−1^ fluorescein (FITC) labelled 500 kDa dextran fraction (Sigma-Aldrich, FD500S) dissolved in injection buffer (50 mM K-HEPES, pH 7.4 (Sigma-Aldrich, H3375), 5% glycerol (Applichem, A0970), 1 mM Mg(OAc)2 (Sigma-Aldrich, M5661)). Microinjection of mitotic cells (Figs. 2a, c, e, 4a, Extended Data Figs. 4a, 6a, c) was performed using injection settings of 130-150 hPa injection pressure, 0.15 to 0.25 s injection time and 30 hPa compensation pressure. Microinjection of G2 interphase cells (Figs. g, I, k, Extended Data Fig. 4c) was performed using injection settings of 120 hPa injection pressure, 0.4 to 0.5 s injection time and 20 hPa compensation pressure.

For injection of nucleosome arrays (Fig. 2m), 1 volume of injection buffer (50 mM K-HEPES, pH 7.4, 25% glycerol buffer) was added to 1 volume unmodified and 1 volume acetylated nucleosome array solution (1.3 μM final concentration for each nucleosome in the injection buffer). Microinjection of mitotic cells was performed using injection settings of 180-190 hPa injection pressure, 0.35 s injection time, 85 hPa compensation pressure).

For injection of tubulin protein, TRITC-labelled tubulin (Cytoskeleton, TL590M) was dissolved in 5%-GPEM (80 mM K-PIPES, pH 6.9 (Sigma-Aldrich, P6757), 1 mM MgCl_2_ (Sigma-Aldrich, 63065), 1 mM EGTA (Merck, 324626)) supplemented with 1 mM GTP (Sigma-Aldrich, G8877) to a concentration of 0.5 mg/ml. Protein was clarified by centrifugation at 4°C for 15 min in a tabletop centrifuge at 21,000 x g. Supernatant was microinjected into mitotic cells using injection settings of 175 hPa injection pressure, 0.25 s, 85 hPa compensation pressure. For microinjection of cells arrested in prometaphase with nocodazole, the G-PEM was additionally supplemented with 100 ng ml^−1^ nocodazole (Sigma-Aldrich, M1404).

For microinjection of GFP surface charge variants (Extended Data Fig. 5a), recombinant mEGFP(−7e) or scGFP(+7e) were dissolved in injection buffer (50 mM K-HEPES, pH 7.4 (Sigma-Aldrich, H3375), 25% glycerol (Applichem, A0970)) to a concentration of 15 μM. Microinjection of mitotic cells was performed using injection settings of 125 hPa injection pressure, 0.2 s injection time, 40 hPa compensation pressure.

For microinjection of charge modified dextran fractions (Extended Data Fig. 5c), a fluorescein (FITC) labelled 4.4 kDa dextran fraction (negative overall charge conferred to dextran by fluorophore charge) (Sigma-Alrich, FD4) or a FITC labelled 4.4 kDa dextran fraction modified with diethylaminoethyl (DEAE) groups conferring overall positive charge (TdB, DD4) were dissolved in injection buffer (50 mM K-HEPES, pH 7.4 (Sigma-Aldrich, H3375), 25% glycerol (Applichem, A0970)) to a concentration of 5 mg ml^−1^. Microinjection of mitotic cells was performed using injection settings of 125 hPa injection pressure, 0.1-15 s injection time, 20 hPa compensation pressure.

### Image analysis

#### DNA congression analysis

To quantify DNA congression to the spindle equator in live cells, the DNA distribution along a line profile (Fig. 1b, 7.06 μm width, 22.5 μm length, Extended Data Fig. 3 c, d, 12.04 μm width, 22.5 μm length) parallel to the pole-to-pole axis was measured. Along each line profile (22.5 μm length in total), the accumulated DNA density in the central 5 μm interval around the 50% distance between the poles (determined by pericentrin stain (Fig. 1a,b) or highest SiR-tubulin staining intensity (Extended Data Fig. 3c,d)) was divided by the total DNA density along the entire profile, after subtraction of the extracellular background fluorescence, with every line profile consisting of an average intensity projection of Z-slices around the pole-to-pole axis (Fig. 1a,b, 2.4 μm range, Extended Data Fig. 3 c, d, 0.75 μm range).

#### Chromatin/DNA compaction analysis

To quantify DNA density, in a central z-section of a mitotic cell (determined by visual inspection based on highest SiR-tubulin staining intensity at the poles (Fig. 1a, c, Extended Data Fig. 3c, e)) the DNA channel was denoised using a Gaussian blur filter (σ=2) and thresholded using the Otsu dark method in Fiji. The resulting binary mask was converted into a selection and the DNA density within this region of interest (ROI) measured. All data points were normalized to the mean of unperturbed control cells.

In STLC-treated cells used for AluI microinjection experiment (Fig. 2a, e, Extended Data Fig. 4c), the DNA density was measured in line profiles. In a single Z-slice, a line profile (3 pixels wide, 2 μm long) through a chromosome parallel to the imaging plane (before AluI-injection) or a chromatin droplet (20 min after AluI-injection) was measured. The mean histone 2B fluorescence intensity in a 200 nm interval around the peak value was measured. Values were normalized to the mean of the non-injected control (Fig. 2b, f) or the non-injected condensin degraded control (Extended Data Fig. 4d).

In cells subjected to G2-mitosis induction experiments (Fig. 2g, i, k, Extended Data Fig. 4c), the YFP channel was denoised using a Gaussian blur filter (σ=2) and thresholded using the Otsu dark method in Fiji. The resulting mask was converted into a ROI and the histone 2B-YFP mean fluorescence recorded within this ROI. Per experiment, the values were normalized to the mean of the G2 measurements.

#### Electron tomography analysis

Analysis of electron tomograms were performed in IMOD/ 3dmod, Version 4.11. Annotation of all structures was performed in the ‘Slicer Window’ view, using running Z average intensity projections to increase contrast (microtubule annotation: projection of 10-20 Z-slices, chromatin annotation: 35-50 Z-slices). To account for loss in image sharpness towards the top and bottom of the recorded tomograms, for all tomograms, only the center section of the recorded volume was analyzed (tomograms with 120 to 150 slices, no annotation in the top and bottom 20 slices). For microtubule annotation, a zoom factor of 0.8 to 1.0 was used. For chromatin annotation, a zoom factor of 0.35-0.45 was used. Assignment of microtubules was based on ultrastructural morphology, and assignment of chromatin boundaries was performed based on local grain size, considering the transition of a large-grained, coarsely interspersed particle containing area (cytosol with ribosomes; typical CV>0.4) to a fine-grained, finely interspersed particle containing area (chromatin with nucleosomes; typical CV<0.3 for control, CV<0.4 for TSA) as the chromosome surface. The electron density as a determining factor of a chromatin surface was only used as a secondary direction since TSA treatment perturbs the chromatin density/morphology and thus decreases annotation accuracy. In average, an annotation landmark was set every 5-10 nm for both microtubules and the chromatin surface. After manual annotation of microtubules and a chromatin surface, a model of a meshed chromatin surface was generated, by averaging 5 consecutive annotated slices to increase surface smoothness. Microtubule segment length was measured within cytoplasm and chromatin-internal regions and normalized to the total volume of the respective domain.

#### Mitotic duration measurements in absence or presence of TSA

Fields of asynchronous cells stably expressing histone 2B-mRFP were imaged for up to 4 h. Timing of nuclear envelope breakdown (NEBD) and anaphase onset was determined by visual inspection of chromatin morphology.

#### FRAP analysis of chromatin mobility

To measure FRAP, raw data measurement, background subtraction, data correction and normalization was performed according to ^60^. The measurement of FRAP ROI and background fluorescence signal was performed directly in ZEN software. Total chromatin fluorescence per time-lapse movie was measured using the ‘Time Series Analyzer v3’ (Author: Balaji J, Dept. of Neurobiology, UCLA).

#### FRET-analysis of AurB-FRET biosensor

To determine the mitotic state of control and AluI-injected cells at the G2 to M transition (Fig. 2g,I,k, Extended data Fig. 4c), CFP, FRET and YFP signals of the histone 2B fused Aurora B-FRET biosensor^37^ were recorded at the same time. To quantify FRET efficiency, a custom Fiji script was used. In average intensity projections of 9 z-slices (1 μm each), a nuclear mask was generated using the YFP fluorescence, after denoising using a Gaussian blur filter (σ=5), by thresholding using the Otsu dark method. For each time point, the background for FRET and CFP channel was measure in an extracellular ROI (square of ~1 by 1 μm). After background subtraction, FRET and CFP signals within the nuclear ROI were denoised using a Gaussian blur filter (σ=5) and a FRET/ CFP ratio calculated. The resulting ratiometric image was depicted using the “Fire” lookup-table in Fiji, in an interval of 0 to 1.4 for FRET/CFP.

#### Macromolecule partitioning relative to chromatin

In images of microinjected nucleosome arrays into metaphase cells (Fig. 2m), the DNA channel was denoised using a Gaussian blur filter (σ=2) and thresholded using the Otsu dark method in Fiji. The chromatin mask was converted into a ROI by using the ‘Analyze Particles’ function in Fiji. For the two injected nucleosome arrays of different colors, the mean fluorescence intensity was measured in the DNA ROI and divided by the fluorescence intensity in the cytosol measured in a circular ROI of ~5 μm diameter, with at least 1 μm distance to the chromosomes.

TRITC-tubulin partition coefficient (Fig. 3a, b) was measured >10 min after microinjection in nocodazole arrested cells. At the surface of a chromosome at the edge of the chromosome cluster, mean fluorescence intensity of TRITC was measured along a line profile (5 pixels width, 1.3 μm length) orthogonal to the chromosome surface. Values 1 μm apart within and outside of chromatin were divided, resulting in the measured equilibrium partition coefficient.

Distribution of different expressed DsRed fusions (Fig. 3c, d) was measured using a custom Fiji script. A chromatin mask was generated using thresholding (Default method, Fiji), after Gaussian blur filtering (σ=2) the chromatin channel. The mask was then shrunk by 0.2 μm and mean fluorescence measured within the mask. Cytoplasmic fluorescence was measured within a 0.5 μm wide ring, generated by extending the mask by 1.5 μm. Background mean fluorescence was measured within a small circular region outside of the cell and subtracted from chromatin and cytoplasmic values. Partitioning coefficients were obtained by dividing chromatin by cytoplasmic fluorescence values. Measurements were performed on the central Z-slice.

Distribution of microinjected differently charged GFPs (Extended Data Fig. 5a, b) and differently modified FITC-labelled dextran fractions (Extended Data Fig. c,d) was measured analogously, along line profiles orthogonal to the metaphase plate at the orthogonal surface of the chromatin mass (5 pixels width, 2.5 μm length). Values 1 μm apart within and outside of chromatin were divided, resulting in the measured equilibrium partition coefficient.

The distribution of soluble TRITC-tubulin dimers (Fig. 3e, f), differently charged GFPs (Extended Data Fig. 5e.f) and differently modified FITC-labelled dextran fractions (Extended Data Fig. g,h) within nucleosome array droplets *in vitro* was measured by determining the mean fluorescence intensity in the droplet in a central section of the droplet, in a small circular region covering ~25% of the droplet area in the respective section. In a large, rectangular buffer ROI without droplets, the mean fluorescence in the buffer was measured and the partition coefficient calculated for each field of droplets.

#### Chromatin droplet distribution analysis in cells

Chromatin distribution in pole-peripheral regions was measured at 36 min after nocodazole washout. The spindle pole position was determined as the z-slice with highest tubulin signal (SiR-tubulin (Fig. 4a) or eGFP-α-tubulin (Extended Data Fig. 4a, c)) and the total histone 2B (Fig. 4a, Extended Data Fig. 6a, c) and eGFP-CENPA fluorescence intensity in circular regions of interest (ROIs) (r=5 μm) in maximum intensity projections of 5 z-slices (1 μm offset between slices) around the central section was subtracted from the total cellular fluorescence in maximum intensity projections (cell outline determined by DIC-channel). Each dataset was normalized to the mean of the total fluorescence intensities of all cells.

#### Microtubule density distribution *in vitro*

After deconvolution in ZEN, Airyscan images of fields of microtubules were denoised using a Gaussian blur filter (σ=2). The background was then removed using the ‘subtract background’ function in Fiji, using a rolling ball radius of 740 pixels. The image was converted into 8-bit, and thresholded using the ‘Auto Local Threshold’ (v1.10.1) plugin in Fiji, with the Phansalkar method and a radius of 100. The resulting binary image was skeletonized using the ‘Skeletonize’ function in Fiji. The resulting skeleton length was measured. The chromatin channel was denoised using a Gaussian blur filter (σ=2) and thresholded using the Otsu method in Fiji. The resulting binary image was transformed into a ROI using the ‘Analyze particles’ function (size range: 5-infinity). Microtubule skeleton lengths were measured within chromatin ROIs and surrounding buffer and the ratio of total segment length in chromatin/buffer calculated per image.

#### Validation of auxin-inducible degradation of Smc4-mAID-HaloTAG in live cells

In images of fields of live cells stained with OregonGreen488 HaloTAG ligand, the total fluorescence intensity in average intensity projection was recorded in Fiji. To normalize the values to the cell count, ratios of HaloTAG fluorescence over SiR-DNA intensity were calculated, and the values normalized to the mean of the Hela WT cells.

#### Acetylated histone fluorescence intensity measurements

After indirect immunofluorescence of the respective histone-acetyl marks, a mask of the DNA channel was generated by thresholding using the Otsu method in Fiji. Next, the obtained binary image was converted to ROIs using the ‘Analyze particles’ function (size range: 5-infinity). Mitotic and interphase cells were differentiated manually based chromatin morphology. Within the obtained ROI per cell, the mean fluorescence intensity of the acetyl-mark was recorded, the extracellular background measured in a rectangular ROI subtracted and a ratio of histone mark over DNA mean intensity generated to account for changes in DNA compaction between the different conditions. For comparison of untreated interphase and mitotic cells, obtained ratios were normalized to the mean of untreated interphase cells (Extended Data Fig. 2a, b). For comparison of untreated and TSA-treated mitotic cells, ratios were normalized to the mean of untreated mitotic cells (Extended Data Fig. 2c, d). For comparison of p300-expressing mitotic cells, ratios were normalized to the mean of the mock-plasmid transfected mitotic cells (Extended Data Fig. 3a, b).

#### Measurement of Cyclin B1 stain

In interphase mitotic cells, determined by absence or presence of cell rounding, in the central section of the recorded Z-stack (7 Z-slices, 1 μm each, central section manually determined), the cell outline in DIC images was used to determine a ROI per cell. The total fluorescence intensity of the CyclinB1 fluorescence within this ROI was recorded in Fiji. The values were normalized to the mean of the CyclinB1 stain in interphase cells.

#### Determination of coefficient of variation of chromatin droplets *in vitro*

In entire fields of chromatin droplets or nucleosome array solution (Extended Data Fig. 4e) (both recorded ~3 μm above the cover glass using laser powers adjusted to the unmodified condition), the mean fluorescence intensity and standard deviation was measured, and the coefficient of variation (CV=α μ^−1^) was calculated.

#### Airyscan processing and deconvolution

Raw images recorded using the Airyscan2 detector using AS-SR or multiplex airyscan modes were processed using ZEN3.3 Blue 2020 software.

#### Fiji version details

The Fiji-integrated distribution/version used for analyses in this study was ImageJ 1.53c, using Java 1.8.0_66 (64-bit), with in part custom ImageJ plugins as indicated.

#### Statistical analyses and data reporting

No statistical methods were used to predetermine sample size. Data were tested for normality and equal variances with Shapiro–Wilk/D’Agostino–Pearson and F-test/Levene’s tests (*α* = 0.05), respectively. The appropriate statistical test was chosen as follows: Data were tested for normality. Unpaired, normally distributed data were tested with a two-tailed *t*-test (in the case of similar variances) or with a two-tailed *t*-test with Welch’s correction (in the case of different variances).

Unpaired, not-normally distributed data were tested with a two-tailed Mann–Whitney test (in the case of similar variances) or with a two-tailed Kolmogorov–Smirnov test (in the case of different variances). Paired, normally distributed data were tested with two-tailed *t*-test and paired, not-normally distributed data were tested with a Wilcoxon matched-pairs signed rank test. When manual annotation was required, blinding precautions were made.

#### Sample numbers

Fig. 1a, b: representative examples and quantification of chromatin fraction at spindle center: Control (*n*=51 cells), ΔCondensin (*n*=65 cells), ΔCondensin+TSA (*n*=34 cells), TSA (*n*=61). Fig. 1c: quantification of DNA density: Control (*n*=31 cells), ΔCondensin (*n*=89 cells), ΔCondensin+TSA (*n*=99 cells), TSA (*n*=74). Fig. 1d,e: representative examples and quantification of microtubule densities: Control (*n*=4 tomograms from 3 different cells), TSA (*n*=4 tomograms from 3 different cells). Fig. 1f: quantification of microtubule segment length in cytoplasm: Control (*n*=4 tomograms from 3 different cells), TSA (*n*=4 tomograms from 3 different cells). Fig. 2a, b: representative example and quantification of 11 cells, 3 ROIs per cell. Fig. 2c, d: representative examples and quantification of 8 cells (Undigested) and 10 cells (AluI-digested). Fig. 2e, f: Representative example and quantification of 11 cells, 3 ROIs per cell. Fig. 2g, h: Representative example and quantification of 13 cells. Fig. 2i, j: Representative example and quantification of 8 cells. Fig. 2k, l: Representative example and quantification of 10 cells. Fig. 2m, n: Representative example and quantification of 28 cells. Fig. 3a: Representative example of an untreated metaphase cell (*n*=27). Fig. 3a, b: Representative examples and quantification of nocodazole treated cells: Control (*n*=46), TSA (*n*=31). Fig. 3c, d: Representative examples and quantification of: DsRed (*n*=26), DsRed(−7e) (*n*=26), DsRed(+9e) (*n*=26). Fig. 3e, f: Representative examples and quantification of: Nocodazole (*n*=94 from 45 fields of droplets), polymerized microtubules (*n*=13 fields of droplets). Fig. 4 a, b: Representative example and quantification of 15 cells. Extended Data Fig. 1a, b: Representative examples and quantification of HeLa WT (*n*=25), HeLa Smc4-mAID-Halo (*n*=25), HeLa Smc4-mAID-Halo + 5-PhIAA (*n*=25). Extended Data Fig. 1d: Representative examples of in total recorded cells: Control (*n*=20), ΔCondensin (*n*=20), ΔCondensin+TSA (*n*=45), TSA (*n*=40). Extended Data Fig. 1e: Quantification of mitotic duration in control (*n*=44) and TSA-treated (*n*=36) cells. Extended Data Fig. 1f, g: Representative examples and quantification of control (*n*=64) and TSA-treated (*n*=110). Extended Data Fig. 2a, b: Representative examples and quantification of interphase (H2B-Ac (*n*=20), H3-Ac (*n*=20), H4K16-Ac (*n*=20)) and mitotic (H2B-Ac (*n*=20), H3-Ac (*n*=20), H4K16-Ac (*n*=20)) cells. Extended Data Fig. 2c, d: Representative examples and quantification of control (see Extended Data Fig. 2b) and TSA treated (H2B-Ac (*n*=20), H3-Ac (*n*=20), H4K16-Ac (*n*=20)). Extended data Fig. 2e, f: Representative examples and quantification of interphase (*n*=60), untreated mitotic (*n*=89), ΔCondensin (*n*=59), ΔCondensin+TSA (*n*=83), TSA (*n*=67). Extended Data Fig. 3a, b: representative examples and quantification of mock (*n*= 20), p300_HAT_ (*n*=26), p300(D1399Y) (*n*=20) transfected cells. Extended Data Fig. 3c, d, e: representative examples and quantification of chromatin fraction at spindle center (d) and chromatin density (e) of mock (*n*=20), ΔCondensin+p300_HAT_ (*n*=20), ΔCondensin+300(D1399Y) (*n*=24) transfected cells. Extended Data Fig. 4a, b: Representative example and quantification of 7 cells, 3 ROIs per cell. Extended Data Fig. 4c, d: Representative example and quantification of 11 cells. Extended Data Fig. 4e, f: Representation examples and quantification of fields of unmodified (AF488: *n*=26, AF594: *n*=25) and acetylated (AF488: *n*=25, AF594: *n*=30) nucleosome arrays. Extended Data Fig. 5a, b: Representative examples and quantification of microinjected mEGFP (*n*=17) and scGFP(*n*=20). Extended Data Fig. 5c, d: Representative examples and quantification of microinjected FITC-dextran (−) (*n*=21) and FITC-DEAE-dextran (+) (*n*=10). Extended Data Fig. 5e, f: Representative examples and quantification of nucleosome array droplets with mEGFP (*n*=69) and scGFP (*n*=73). Extended Data Fig. 5g, h: Representative examples and quantification of nucleosome array droplets with FITC-dextran (−) (*n*=69) and FITC-DEAE-dextran (+) (*n*=57). Extended Data Fig. 6 a, b: Representative example and quantification of 13 cells. Extended Data Fig. c, d: Representative example and quantification of 16 cells. All experiments in this study were performed in at least 2 biological replicates.

## Acknowledgments

The authors thank the IMBA/IMP/GMI BioOptics and Molecular Biology Service and the VBCF Electron Microscopy and Protein Technologies facilities for technical support. The authors thank Iain Patten, Alexey Khodjakov, Helder Maiato, and Carsten Janke for comments on the manuscript. The authors thank Paul Batty for assistance with cell line generation and Angela M. Rodrigues Viana for advice and reagents for genome engineering. Research in the laboratory of D.W.G. is supported by the Austrian Academy of Sciences, the Austrian Science Fund (FWF; Doktoratskolleg “Chromosome Dynamics” DK W1238), and the Vienna Science and Technology Fund (WWTF; projects LS17-003 and LS19-001). Research in the laboratory of S.O. is supported by the Vienna Science and Technology Fund (WWTF; project LS19-001). Research in the laboratory of M.K.R. is supported by the Howard Hughes Medical Institute, a Paul G. Allen Frontiers Distinguished Investigator Award (to M.K.R.) and grants from the NIH (F32GM129925 to B.A.G.) and the Welch Foundation (I-1544 to M.K.R.). M.W.G.S. and M.P. have received a PhD fellowship from the Boehringer Ingelheim Fonds.

## Author contributions

D.W.G. and M.W.G.S. conceived the project, with input from B.A.G. and M.K.R.. M.W.G.S. designed, performed, and analyzed all experiments, except those shown in Fig. 1 d-f (M.W.G.S. together with S.O), Figs. ED 4a, b, ED 5e-h (M.W.G.S. together with C.B.), Figs. ED 1 a, b, e, ED 2 a-d, ED 3 a-c (M.F.D.S.), Fig. 3 c, d (M.P.). B.A.G. and L.K.D. generated nucleosome arrays and developed *in vitro* chromatin condensate assays. C.C.H.L. developed image analysis procedures. D.W.G and M.K.R acquired funding and supervised the project. D.W.G and M.W.G.S wrote the manuscript, with input from all authors.

---

**Extended Data Figure 1.**
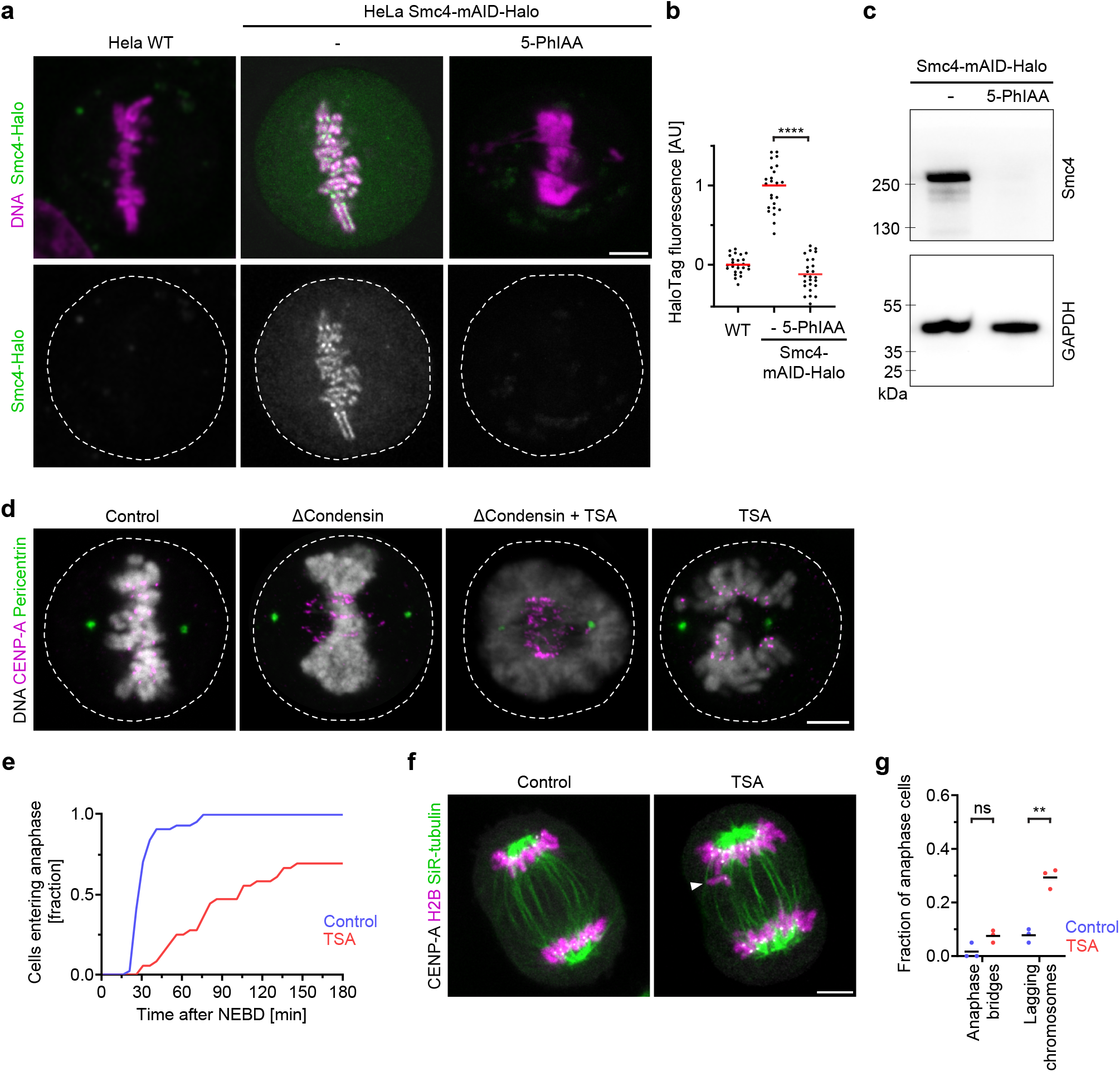
Characterization of HeLa Smc4-mAID-HaloTAG cells and effect of TSA on chromosome segregation. **a-c**, Smc4 expression analysis. **a**, HeLa cells with homozygous Smc4-mAID-Halo alleles and stably expressing OsTIR(F74G) were analyzed with or without treatment of 5-Phenylindole-3-eacetic acid (5-PhIAA) for 3 hours and stained with OregonGreen-488 HaloTAG stain; HeLa wildtype (WT) cells serve as control to measure fluorescence background. DNA was stained with Hoechst 33342. **b**, Quantification of HaloTAG fluorescence in live mitotic cells. *n*=25 for WT, *n*=25 for Smc4-mAID-Halo without 5-PhIAA, *n*=25 for Smc4-mAID-Halo with 5-PhIAA. Bars indicate mean; significance was tested by a two-tailed Welch *t*-test (Smc4-mAID-Halo with 5-PhIAA, *P*<10^−15^). **c,** Immunoblot analysis of Smc4-mAID-Halo cells with or without 3h 5-PhIAA treatment. **d**, Immunofluorescence analysis of centromeres and spindle poles in mitotic Smc4-mAID-Halo cells after 3 h degradation of Smc4 (ΔCondensin), 3 h treatment with TSA as indicated. Projection of 7 Z-sections. **e**. Mitotic progression analysis by time-lapse microscopy of HeLa cells expressing H2B-mRFP, in untreated control and TSA-treated cells. *n*=44 for control from 5 biological replicates, *n*=36 for TSA from 4 biological replicates. **f**. Chromosome missegregation analysis by live Airyscan imaging of HeLa cells expressing H2B-mCherry and mEGFP-CENP-A and stained with SiR-tubulin. Representative images of untreated controls or 3 h TSA-treated anaphase cells. Single Airyscan Z‑slices. **g**. Quantification of chromosome missegregation as illustrated in **f**. Dots indicate biological replicates, bars indicate mean. *n*=64 cells for control, *n*=110 for TSA. Significance was tested by an unpaired, two-tailed *t*-test (anaphase bridges, *P*=0.052, lagging chromosomes. *P*=1.209×10^−3^). Scale bars, 5 μm.

**Extended Data Figure 2.**
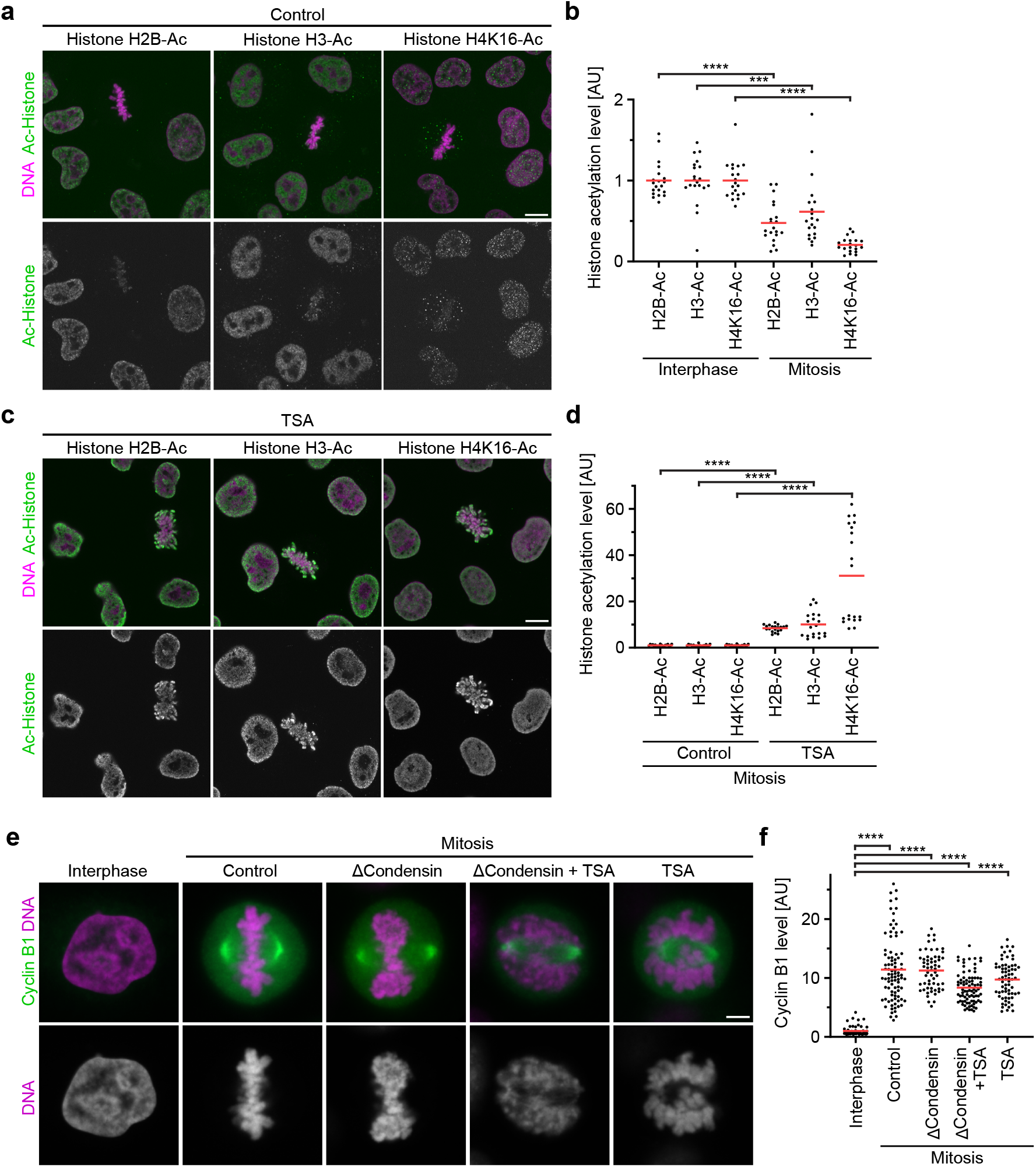
Analysis of histone acetylation and cell cycle states. **a**,**b**, Immunofluorescence analysis of histone acetylation in interphase and mitosis. **a**, HeLa cells were fixed and stained with antibodies against different acetylated histones for immunofluorescence as indicated. **b**, Quantification of histone acetylation by immunofluorescence as in **a**, by the ratio of antibody fluorescence to DNA reference staining by Hoechst 33342. For each acetylated histone, all data points were normalized to the mean of interphase cells. *n*=20 for H2B-Ac, Interphase, *n*=20 for H3-Ac, Interphase, *n*=20 for H4K16-Ac, Interphase, *n*=20 for H2B-Ac, Mitosis, *n*=20 for H3-Ac, Mitosis, *n*=20 for H4K6-Ac, Mitosis. Bars indicate mean; significance was tested by a two-tailed Welch *t*-test (H2B-Ac, Mitosis, *P*=2.572×10^−8^), or a two-tailed Mann-Whitney test (H3-Ac, Mitosis, *P*=4.72×10^−4^), or a Kolmogorov-Smirnov test (H4K16Ac, Mitosis, *P*=4.122x). **c**,**d**, Histone acetylation in mitotic cells 3 h after TSA treatment. **c**, HeLa cells were treated with TSA for 3 h, fixed, and histone acetylation analyzed by immunofluorescence as in **a**. **d**, Quantification of histone acetylation as in b, comparing mitotic cells 3 h after TSA treatment with untreated control cells. For each acetylated histone antibody, all data points were normalized to the mean of control metaphase cells. *n*=20 for H2B-Ac, control, *n*=20 for H3-Ac, control, *n*=20 for H4K16-Ac, control, *n*=20 for H2B-Ac, TSA, *n*=20 for H3-Ac, TSA, *n*=20 for H4K16-Ac, TSA. Bars indicate mean; significance was tested by a two-tailed Welch *t*-test (H2B-Ac, TSA, *P*<10^−15^; H3-Ac, TSA, *P*=4.767×10^−7^) or a two-tailed Mann-Whitney test (H4K16-Ac, TSA, *P*=1.451×10^−11^). **e**, **f**, Analysis of cell cycle state by immunofluorescence staining against Cyclin B1. HeLa cells with homozygously mAID-tagged SMC4 were treated with 5-PhIAA to deplete condensin (ΔCondensin) or with TSA to suppress mitotic histone deacetylation as indicated. DNA was stained with Hoechst 33342. Classification of interphase and mitotic cell is based on overall cell shape and spindle morphology. **f**, Quantification of Cyclin B1 fluorescence in cells as in **e**. Data normalized to the mean of untreated interphase cells. *n*=60 (interphase), *n*=89 (mitotic), *n*=59 (ΔCondensin), *n*=83 (ΔCondensin+TSA), *n*=67 (TSA) cells. Bars indicate mean; significance was tested by a two-tailed Mann-Whitney test (CTRL, *P*<10^−15^); ΔCondensin, *P*<10^−15;^ ΔCondensin+TSA, *P*<10^−15^; TSA, *P*<10^−15^). Scale bars, **a**,**c**, 10 μm, **e**, 5 μm.

**Extended Data Figure 3.**
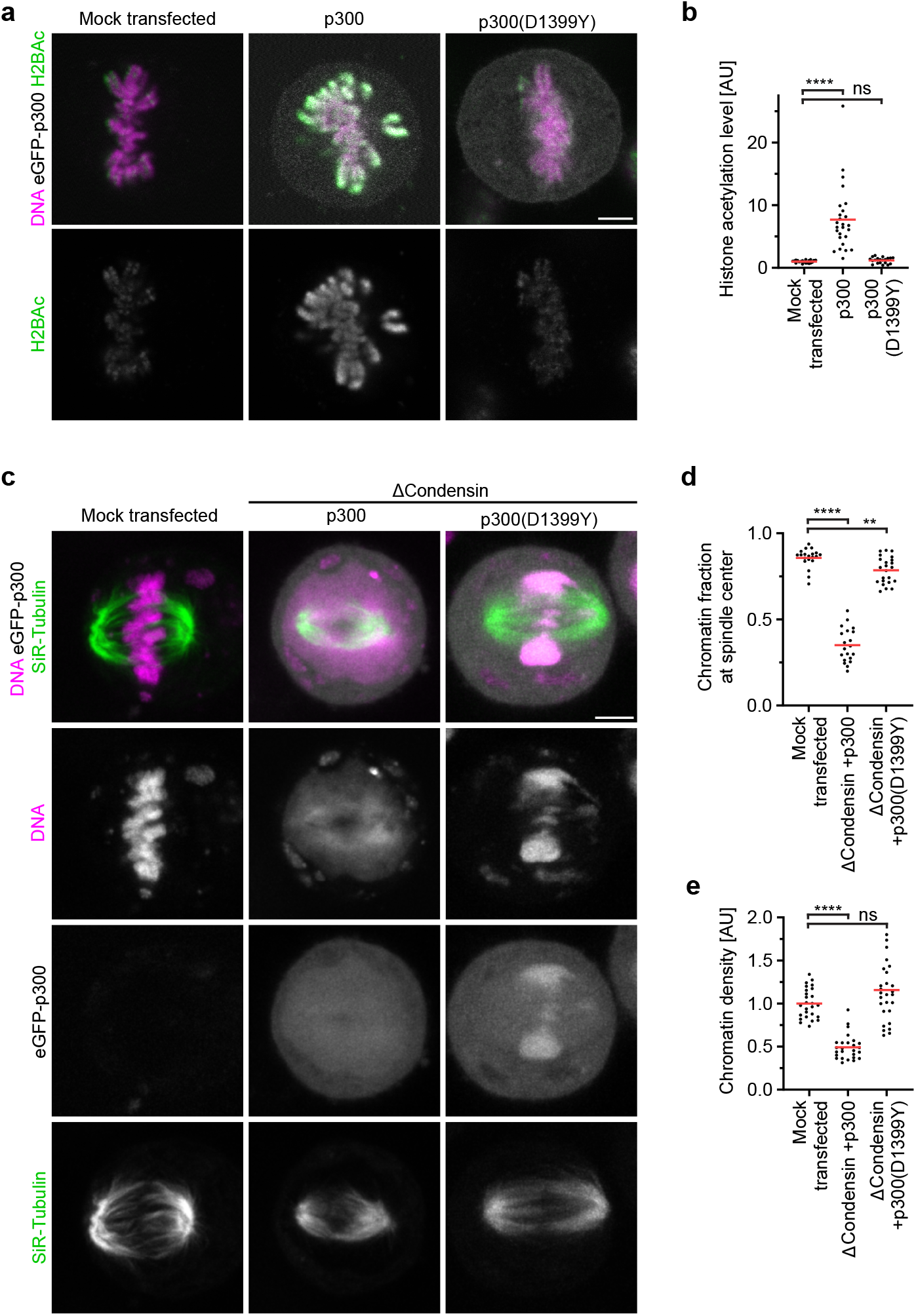
Histone acetylation and chromosome organization in cells overexpressing eGFP-p300. **a**, **b**, Immunofluorescence analysis of histone acetylation in mitotic cells overexpressing p300 histone acetyltransferase or catalytically dead p300(D1399Y). **a**, Cells were transfected with a plasmid coding for eGFP-p300 or eGFP-p300 (D1399Y) and fixed after 48 h. Histone 2B acetylation was analyzed by immunofluorescence. DNA was stained with Hoechst 3342. **b**, Quantification of histone acetylation in metaphase cells as in **a**. Data points were normalized to the mean of mock-transfected mitotic cells. *n*=20 for mock-transfected, *n*=26 for p300, *n*=20 for p300 (D1399Y). Bars indicate mean; significance was tested by a two-tailed Mann-Whitney test (p300, *P*=3.57×10^−13^) or a two-tailed Welch *t*-test (p300(D1399Y), *P*=0.138). **c**-**e**, Analysis of chromatin density and congression to spindle center in cells after SMC4-AID-Halo degradation (ΔCondensin) and overexpression of p300 or catalytically dead p300(D1399Y). **c**, Cells transfected with a plasmid coding for eGFP-p300 or eGFP-p300 (D1399Y) for 48 hours were stained with Hoechst 33342 and SiR-Tubulin and mitotic cells with bipolar spindles imaged live. Projection of 5 z-sections. **d**, Quantification of chromosome congression by the fraction of chromatin localizing to the central spindle region. *n*=20 for mock-transfected, *n*=24 for ΔCondensin+p300, *n*=20 for ΔCondensin+p300(D1399Y). Bars indicate mean; significance was tested by a Kolmogorov-Smirnov test (ΔCondensin+p300, *P*=4.122×10^−9^) or a two-tailed Mann-Whitney test (ΔCondensin+p300(D1399Y), *P*=1.72×10^−3^). **e**, Quantification of chromatin density in cells treated as in **c**. *n*=20 for mock-transfected, *n*=24 for ΔCondensin+p300, *n*=20 for ΔCondensin+p300(D1399Y). Bars indicate mean; significance was tested by a two-tailed Mann-Whitney test (ΔCondensin+p300, *P*=7.86×10^−13^) or a two-tailed Welch *t*-test (ΔCondensin+p300(D1399Y), *P*=0.051). Scale bars, 5 μm.

**Extended Data Figure 4.**
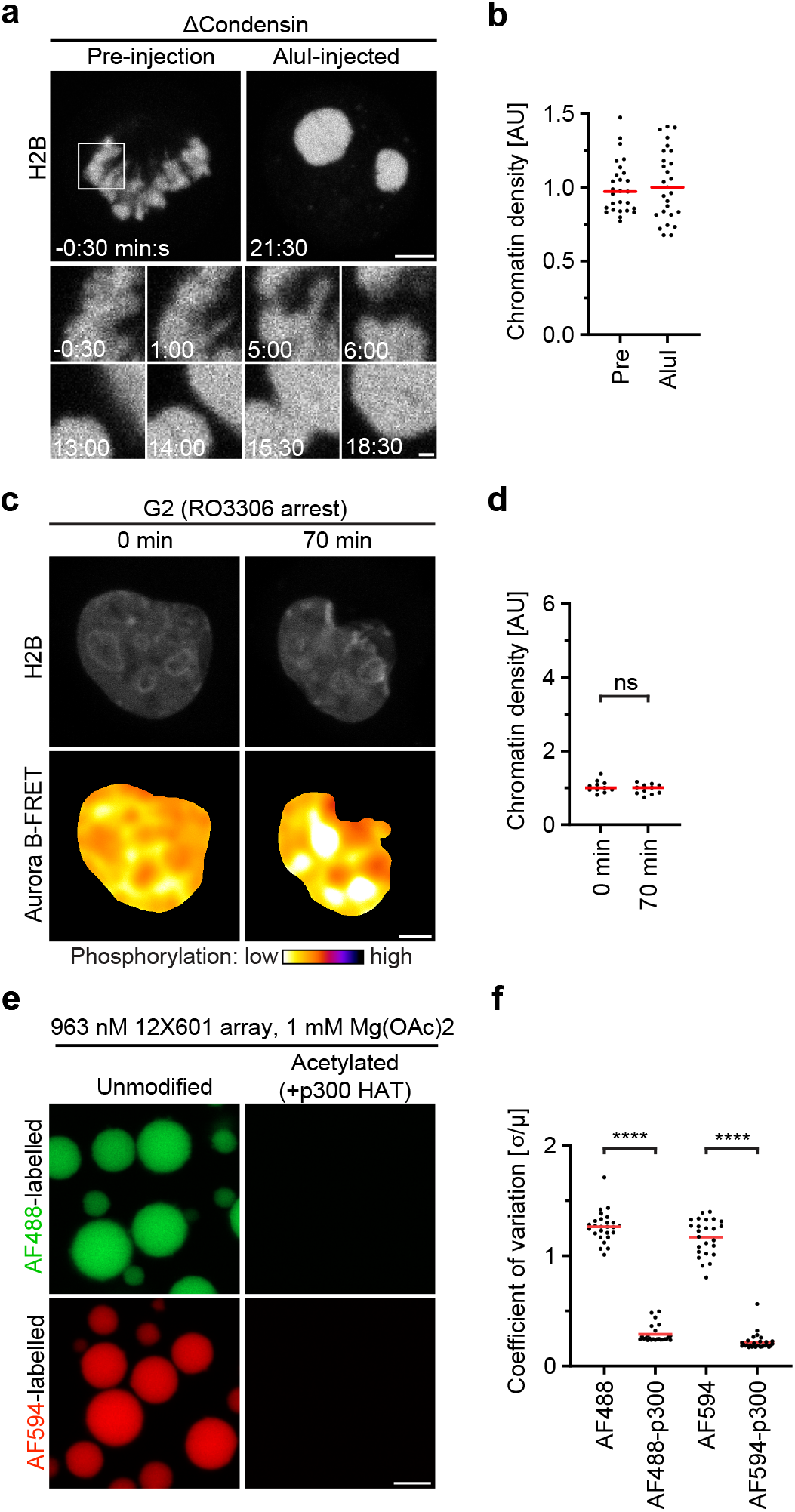
Phase separation behavior of chromatin fragments. **a**, **b**, Condensin is not required for liquid phase separation of chromatin fragments in mitotic cells. **a**. Smc4-AID HeLa cells expressing H2B-mCherry were treated 3 h with 5-PhIAA to deplete condensin and a mitotic cell was then injected with AluI (t=0 min) and imaged by time-lapse microscopy. **b**, Quantification of chromatin density before and after injection of AluI, normalized to the mean of untreated pre-injection cells. *n*=7 cells, 3 ROIs each. Bars indicate mean. **c**, **d**, Chromatin fragments do not undergo liquid phase separation in G2 cells. **c**, HeLa cell expressing Aurora B-FRET biosensor was synchronized to G2 by RO3306 and microinjected in the nucleus with AluI. G2 state was retained in presence of RO3306 as indicated by FRET signal. t=0 minutes refers to the first timepoint of the recorded time-lapse. **d**, Quantification of chromatin density in cells as in **c**, normalized to the mean of t=0 min. *n*=11 cells. Bars indicate mean; significance tested by a paired, two-tailed *t*-test (*P*=0.091). **e**, **f**, In vitro liquid-liquid phase separation behavior of unmodified or acetylated nucleosome arrays. **e**, 12X601 Nucleosome arrays labelled with fluorophores as indicated were treated with recombinant p300 histone acetyltransferase or no enzyme *in vitro* and then subjected to identical phase separation buffers for 30 minutes. **f**, Quantification of nucleosome array self-association by coefficient of variation (CV= σ/μ) in images as in **e**. *n*=26 for AlexaFlour488 array (AF488), *n*=25 for acetylated AlexaFluor488 array (AF488-p300), *n*=25 for AlexaFluor594 array (AF594), *n*=30 for acetylated AlexaFluor594 array (AF594-p300). Bars indicate mean; significance tested by a Kolmogorov-Smirnov test (AF488-p300, *P*=1.701×10^‑11^; AF594-p300, *P*=2.862×10^−12^). Scale bars, 5 μm, **a**, insert 1 μm.

**Extended Data Figure 5.**
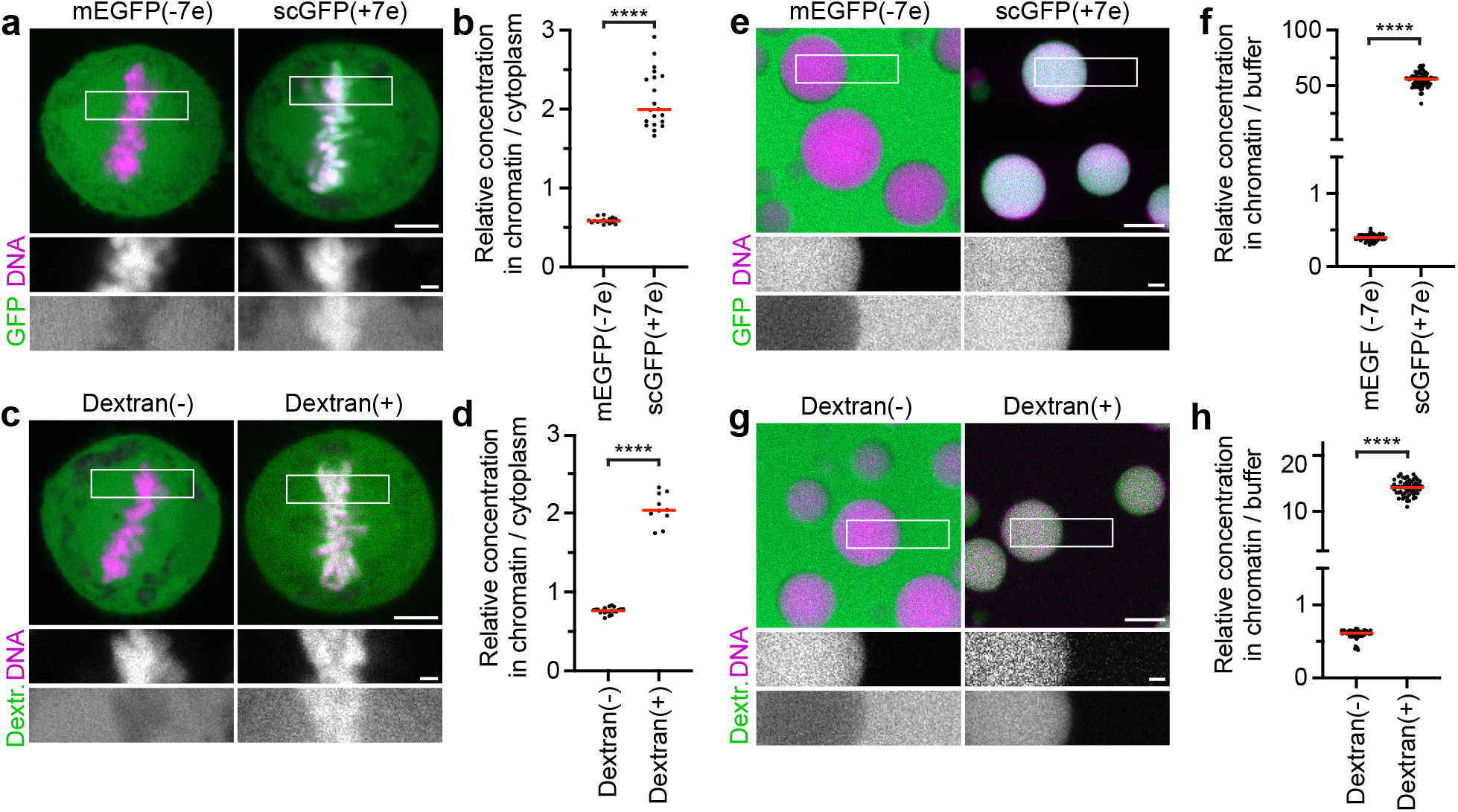
Partitioning of proteins and dextrans relative to chromatin depends on electrical charge. **a**, **b**, Partitioning of GFP surface charge variants relative to chromatin in metaphase cells. **a**, Live metaphase cells after injection of recombinant mEGFP(‑7e) or scGFP(+7e). DNA stained with Hoechst 33342. **b**, Quantification of GFP fluorescence in chromatin relative to cytoplasm. *n*=17 for mEGFP(−7e), *n*=20 for scGFP(+7e). Bars indicate mean; significance tested by a two-tailed Welch *t*-test (*P*=5.6×10^−14^). **c**, **d**, Partitioning of charge-modified fluorescent dextrans relative to chromatin in metaphase cells. **c**, Live metaphase cells after injection of negatively or positively charged 4.4 kDa dextrans. DNA stained with Hoechst 33342. **d**, Quantification of dextran fluorescence in chromatin relative to cytoplasm. *n*=21 for 4.4 kDa dextran (−), *n*=10 for 4.4 kDa dextran(+). Bars indicate mean; significance tested by a two-tailed Welch *t*-test (*P*=4.553×10^−9^). **e**, **f**, Partitioning of GFP surface charge variants relative to liquid droplets of nucleosome arrays *in vitro*. **e**, Liquid chromatin droplets were formed *in vitro* by exposing 12X601 nucleosome arrays to phase separation buffer. GFP charge variants were added for 10 minutes and then imaged. DNA was stained with Hoechst 33342. **f**, Quantification of GFP fluorescence in chromatin relative to buffer. *n*=69 for mEGFP(−7e), *n*=73 for scGFP(+7e). Bars indicate mean; significance tested by a two-tailed Welch *t*-test (*P*<10^−15^). **g**, **h**, Partitioning of charge modified dextrans relative to liquid droplets of nucleosome arrays *in vitro*. **g**, Liquid chromatin droplets were formed were formed as in **e**. Charge modified 4.4 kDa dextrans were added for 10 minutes and then imaged. DNA was stained with Hoechst 33342. **h**, Quantification of dextran fluorescence in chromatin relative to buffer. *n*=69 for 4.4 kDa dextran(−), *n*=57 for 4.4 kDa dextran(+). Bars indicate mean; significance tested by a two-tailed Mann-Whitney test (*P*<10^‑15^). Scale bars, 5 μm, inserts 1 μm.

**Extended Data Figure 6.**
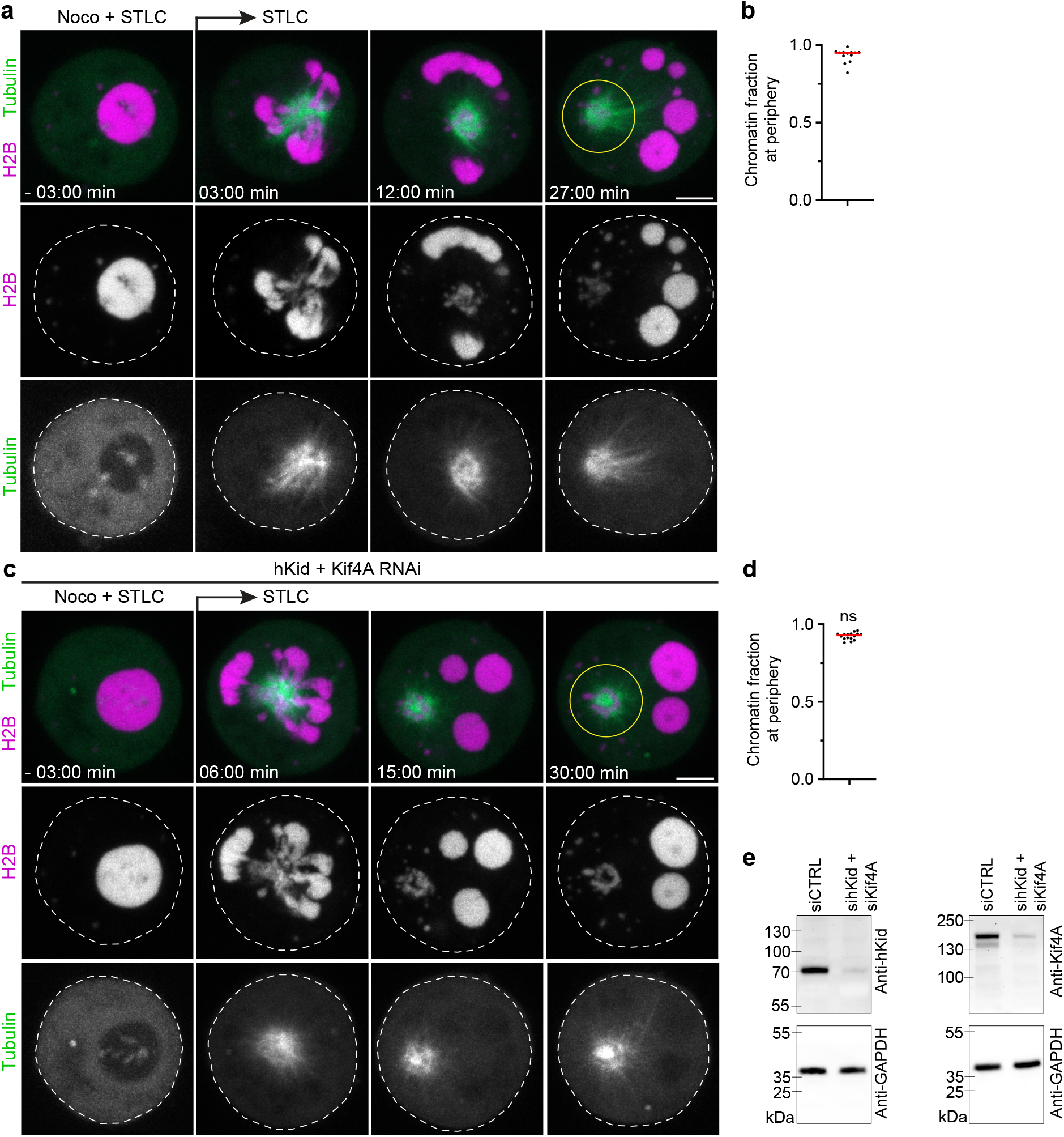
Microtubules push liquified chromatin away from the spindle pole independently of hKid and Kif4A. **a**, Time-lapse microscopy of liquified chromatin during chemically-induced assembly of monopolar spindles. Live mitotic HeLa cells expressing H2B-mCherry and eGFP-α-tubulin were treated with nocodazole and STLC and then injected with AluI. Nocodazole was removed at t=0 min during time-lapse imaging. Projection of 4 z-sections. **b**, Quantification of chromatin localization at the cell periphery relative to the region around the spindle monopole at = 36 min. *n*=13 cells. **c,** Time-lapse microscopy of liquified chromatin during spindle assembly as in **a** for cells depleted of Kid and Kif4a by RNAi. **d,** Quantification as in **b** for hKid/Kif4a-RNAi cells. *n*=16 cells. Bar indicates mean; significance tested by a two-tailed Mann-Whitney test (*P*=0.215) **e,** Validation of RNAi efficiency by Western Blotting. Samples were collected 30 h after transfection of siRNAs targeting hKid and Kif4a and probed by antibodies as indicated. Scale bars, 5 μm.

## References

1. Batty, P. & Gerlich, D. W. Mitotic Chromosome Mechanics: How Cells Segregate Their Genome. Trends in Cell Biology 29, 717–726 (2019).

2. Kschonsak, M. & Haering, C. H. Shaping mitotic chromosomes: From classical concepts to molecular mechanisms. BioEssays : news and reviews in molecular, cellular and developmental biology 37, 755–766 (2015).

3. Ohta, S., Wood, L., Bukowski-Wills, J.-C., Rappsilber, J. & Earnshaw, W. C. Building mitotic chromosomes. Current Opinion In Cell Biology 23, 114–121 (2011).

4. Hirano, T. & Kobayashi, R. Condensins, Chromosome Condensation Protein Complexes Containing XCAP-C, XCAP-E and a Xenopus Homolog of the Drosophila Barren Protein. Cell 89, 511–521 (1997).

5. Shintomi, K., Takahashi, T. S. & Hirano, T. Reconstitution of mitotic chromatids with a minimum set of purified factors. Nature Cell Biology 17, 1014–1023 (2015).

6. Ganji, M. et al. Real-time imaging of DNA loop extrusion by condensin. Science (New York, NY) eaar7831 (2018).

7. Gibcus, J. H. et al. A pathway for mitotic chromosome formation. Science 359, eaao6135 (2018).

8. Yatskevich, S., Rhodes, J. & Nasmyth, K. Organization of Chromosomal DNA by SMC Complexes. Annual Review of Genetics 53, annurev-genet-112618-043633 (2019).

9. Cimini, D., Mattiuzzo, M., Torosantucci, L. & Degrassi, F. Histone hyperacetylation in mitosis prevents sister chromatid separation and produces chromosome segregation defects. Molecular biology of the cell 14, 3821–3833 (2003).

10. Kruhlak, M. J. et al. Regulation of global acetylation in mitosis through loss of histone acetyltransferases and deacetylases from chromatin. The Journal of biological chemistry 276, 38307–38319 (2001).

11. Wilkins, B. J. et al. A cascade of histone modifications induces chromatin condensation in mitosis. Science (New York, NY) 343, 77–80 (2014).

12. Zhiteneva, A. et al. Mitotic post-translational modifications of histones promote chromatin compaction in vitro. Open biology 7, 170076 (2017).

13. Gibson, B. A. et al. Organization of Chromatin by Intrinsic and Regulated Phase Separation. Cell 179, 470–484.e21 (2019).

14. Gerlich, D., Hirota, T., Koch, B., Peters, J. M. & Ellenberg, J. Condensin I stabilizes chromosomes mechanically through a dynamic interaction in live cells. Current Biology 16, 333–344 (2006).

15. Houlard, M. et al. Condensin confers the longitudinal rigidity of chromosomes. Nature Cell Biology 17, 771–781 (2015).

16. Sun, M., Biggs, R., Hornick, J. & Marko, J. F. Condensin controls mitotic chromosome stiffness and stability without forming a structurally contiguous scaffold. Chromosome research : an international journal on the molecular, supramolecular and evolutionary aspects of chromosome biology 26, 277–295 (2018).

17. Oriola, D., Needleman, D. J. & Brugués, J. The Physics of the Metaphase Spindle. Annual Reviews of Biophysics 47, 655–673 (2018).

18. Monda, J. K. & Cheeseman, I. M. The kinetochore-microtubule interface at a glance. Journal Of Cell Science 131, jcs214577 (2018).

19. Rieder, C. L., Davison, E. A., Jensen, L. C., Cassimeris, L. & Salmon, E. D. Oscillatory movements of monooriented chromosomes and their position relative to the spindle pole result from the ejection properties of the aster and half-spindle. The Journal of cell biology 103, 581–591 (1986).

20. Ault, J. G., DeMarco, A. J., Salmon, E. D. & Rieder, C. L. Studies on the ejection properties of asters: astral microtubule turnover influences the oscillatory behavior and positioning of mono-oriented chromosomes. J Cell Sci 99 (Pt 4), 701–710 (1991).

21. Rieder, C. L. & Salmon, E. D. Motile kinetochores and polar ejection forces dictate chromosome position on the vertebrate mitotic spindle. The Journal of cell biology 124, 223–233 (1994).

22. Maiato, H., Gomes, A. M., Sousa, F. & Barisic, M. Mechanisms of Chromosome Congression during Mitosis. Biology 6, 13 (2017).

23. Barisic, M., Aguiar, P., Geley, S. & Maiato, H. Kinetochore motors drive congression of peripheral polar chromosomes by overcoming random arm-ejection forces. Nature Cell Biology 16, 1249–1256 (2014).

24. Yesbolatova, A. et al. The auxin-inducible degron 2 technology provides sharp degradation control in yeast, mammalian cells, and mice. Nature Communications 11, 5701 (2020).

25. Lukinavičius, G. et al. Fluorogenic probes for live-cell imaging of the cytoskeleton. Nature Methods 11, 731–733 (2014).

26. Ginno, P. A., Burger, L., Seebacher, J., Iesmantavicius, V. & Schübeler, D. Cell cycle-resolved chromatin proteomics reveals the extent of mitotic preservation of the genomic regulatory landscape. Nature communications 9, 4012–4048 (2018).

27. Yoshida, M., Kijima, M., Akita, M. & Beppu, T. Potent and specific inhibition of mammalian histone deacetylase both in vivo and in vitro by trichostatin A. The Journal of biological chemistry 265, 17174–17179 (1990).

28. Ogryzko, V. V., Schiltz, R. L., Russanova, V., Howard, B. H. & Nakatani, Y. The Transcriptional Coactivators p300 and CBP Are Histone Acetyltransferases. Cell 87, 953–959 (1996).

29. Hudson, D. F., Vagnarelli, P., Gassmann, R. & Earnshaw, W. C. Condensin is required for nonhistone protein assembly and structural integrity of vertebrate mitotic chromosomes. Developmental cell 5, 323–336 (2003).

30. Samejima, K. et al. Functional analysis after rapid degradation of condensins and 3D-EM reveals chromatin volume is uncoupled from chromosome architecture in mitosis. Journal Of Cell Science 131, jcs210187 (2018).

31. Marko, J. F. & Siggia, E. D. Polymer models of meiotic and mitotic chromosomes. Molecular biology of the cell 8, 2217–2231 (1997).

32. Poirier, M. G. & Marko, J. F. Mitotic chromosomes are chromatin networks without a mechanically contiguous protein scaffold. Proceedings of the National Academy of Sciences of the United States of America 99, 15393–15397 (2002).

33. Poirier, M. G., Monhait, T. & Marko, J. F. Reversible hypercondensation and decondensation of mitotic chromosomes studied using combined chemical-micromechanical techniques. Journal of cellular biochemistry 85, 422–434 (2002).

34. Cuylen, S., Metz, J. & Haering, C. H. Condensin structures chromosomal DNA through topological links. Nature structural & molecular biology 18, 894–901 (2011).

35. Biggs, R., Liu, P. Z., Stephens, A. D. & Marko, J. F. Effects of altering histone posttranslational modifications on mitotic chromosome structure and mechanics. Molecular biology of the cell 30, 820–827 (2019).

36. Rubinstein, M. & Colby, R. H. Polymer Physics. (Oxford University Press, 2003).

37. Fuller, B. G. et al. Midzone activation of aurora B in anaphase produces an intracellular phosphorylation gradient. Nature 453, 1132–1136 (2008).

38. Vassilev, L. T. et al. Selective small-molecule inhibitor reveals critical mitotic functions of human CDK1. Proceedings of the National Academy of Sciences of the United States of America 103, 10660–10665 (2006).

39. Mershin, A., Kolomenski, A. A., Schuessler, H. A. & Nanopoulos, D. V. Tubulin dipole moment, dielectric constant and quantum behavior: computer simulations, experimental results and suggestions. Biosystems 77, 73–85 (2004).

40. Wall, M. A., Socolich, M. & Ranganathan, R. The structural basis for red fluorescence in the tetrameric GFP homolog DsRed. Nature Structural Biology 7, 1133–1138 (2000).

41. Gebala, M., Johnson, S. L., Narlikar, G. J. & Herschlag, D. Ion counting demonstrates a high electrostatic field generated by the nucleosome. eLife 8, (2019).

42. McNaughton, B. R., Cronican, J. J., Thompson, D. B. & Liu, D. R. Mammalian cell penetration, siRNA transfection, and DNA transfection by supercharged proteins. Proceedings of the National Academy of Sciences 106, 6111–6116 (2009).

43. Dogterom, M. & Yurke, B. Measurement of the Force-Velocity Relation for Growing Microtubules. Science 278, 856–860 (1997).

44. Skoufias, D. A. et al. S-Trityl-L-cysteine Is a Reversible, Tight Binding Inhibitor of the Human Kinesin Eg5 That Specifically Blocks Mitotic Progression*,. Journal of Biological Chemistry 281, 17559–17569 (2006).

45. Levesque, A. A. & Compton, D. A. The chromokinesin Kid is necessary for chromosome arm orientation and oscillation, but not congression, on mitotic spindles. The Journal of cell biology 154, 1135–1146 (2001).

46. Brouhard, G. J. & Hunt, A. J. Microtubule movements on the arms of mitotic chromosomes: Polar ejection forces quantified in vitro. PNAS 102, 13903–13908 (2005).

47. Wandke, C. et al. Human chromokinesins promote chromosome congression and spindle microtubule dynamics during mitosis. The Journal of cell biology 198, 847–863 (2012).

48. Almeida, A. C. & Maiato, H. Chromokinesins. Curr Biol 28, R1131–R1135 (2018).

49. Walther, N. et al. A quantitative map of human Condensins provides new insights into mitotic chromosome architecture. The Journal of cell biology 217, 2309–2328 (2018).

50. Woodruff, J. B. et al. The Centrosome Is a Selective Condensate that Nucleates Microtubules by Concentrating Tubulin. Cell 169, 1066–1077.e10 (2017).

## Methods references

51. Schmitz, M. H. A. et al. Live-cell imaging RNAi screen identifies PP2A-B55α and importin-β 21 as key mitotic exit regulators in human cells. Nature Cell Biology 12, 886–893 (2010).

52. Samwer, M. et al. DNA Cross-Bridging Shapes a Single Nucleus from a Set of Mitotic Chromosomes. Cell 170, 956–972.e23 (2017).

53. Gutschner, T., Haemmerle, M., Genovese, G., Draetta, G. F. & Chin, L. Post-translational Regulation of Cas9 during G1 Enhances Homology-Directed Repair. Cell Reports 14, 1555–1566 (2016).

54. Li, S., Prasanna, X., Salo, V. T., Vattulainen, I. & Ikonen, E. An efficient auxin-inducible degron system with low basal degradation in human cells. Nat Methods 16, 866–869 (2019).

55. Thompson, D. B., Cronican, J. J. & Liu, D. R. Engineering and identifying supercharged proteins for macromolecule delivery into mammalian cells. Methods in enzymology 503, 293–319 (2012).

56. Dixit, R. & Ross, J. L. Chapter 27 - Studying Plus-End Tracking at Single Molecule Resolution Using TIRF Microscopy. in Methods in Cell Biology (eds. Wilson, L. & Correia, J. J.) vol. 95 543–554 (Academic Press, 2010).

57. Field, C. M., Pelletier, J. F. & Mitchison, T. J. Chapter 24 - Xenopus extract approaches to studying microtubule organization and signaling in cytokinesis. in Methods in Cell Biology (ed. Echard, A.) vol. 137 395–435 (Academic Press, 2017).

58. Mastronarde, D. N. Automated electron microscope tomography using robust prediction of specimen movements. Journal of Structural Biology 152, 36–51 (2005).

59. Kremer, J. R., Mastronarde, D. N. & McIntosh, J. R. Computer Visualization of Three-Dimensional Image Data Using IMOD. Journal of Structural Biology 116, 71–76 (1996).

60. Bancaud, A., Huet, S., Rabut, G. & Ellenberg, J. Fluorescence perturbation techniques to study mobility and molecular dynamics of proteins in live cells: FRAP, photoactivation, photoconversion, and FLIP. Cold Spring Harbor protocols 2010, pdb.top90 (2010).

